# Transcriptional control of visual neural circuit development by GS homeobox 1

**DOI:** 10.1101/2022.12.30.522239

**Authors:** Alexandra Rose Schmidt, Rebekah Shephard, Regina L Patrick, Sadie A Bergeron

**Author notes:** <u>Correspondence</u>: Sadie A Bergeron, PhD, West Virginia University Departments of Biology and Neuroscience.

## Abstract

As essential components of gene expression networks, transcription factors regulate neural circuit assembly. *GS homeobox 1 (gsx1*) is expressed in the developing visual system; however, no studies have examined its role in visual system formation. In zebrafish, retinal ganglion cell (RGC) axons terminate in ten arborization fields (AFs) in the optic tectum (TeO) and pretectum (Pr). Pretectal AFs (AF1-AF9) mediate distinct and essential visual behaviors, yet we understand less about their development compared to AF10 in the TeO. Using *gsx1* zebrafish mutants, immunohistochemistry, and transgenic lines, we observed that *gsx1* is required for vesicular glutamate transporter, *slc17a6b*, expression in the Pr, but not overall neuron number. *gsx1* mutants have normal eye morphology, yet exhibit impaired vision and a significantly reduced volume of RGC axons innervating the Pr and TeO, including loss of AF7. Consistent with this, prey capture is reduced in *gsx1* mutants. Timed laser ablation of *slc17a6b-positive* neurons reveals that they aide directly in AF7 formation. This work is the first to implicate *gsx1* in establishing cell identity and functional neural circuits in the visual system.

**SUMMARY STATEMENT:** This is the first study in any vertebrate model to establish a requirement for the homeobox transcription factor encoding gene, *gsx1*, in visual neural circuit formation and function.

## INTRODUCTION

The visual system eye-to-brain circuit is a classic and extensively studied model to uncover neural circuit connectivity mechanisms that dictate innate and complex visually mediated behaviors (Bollmann, 2019; Guido, 2018; Harada et al., 2007). In vertebrates, visual information is transmitted to the brain via retinal ganglion cell (RGC) axons. Once RGC axons exit the eye, well-known molecular cues laid out by forebrain and retinorecipient regions including erythropoietin-producing hepatocellular (Eph) receptors and ephrin ligands (Kita et al., 2015a; Nikolov et al., 2013) and Robo/Slit (Campbell et al., 2007; Fricke et al., 2001; Wyatt et al., 2010; Xiao et al., 2011, 2005) guide axons by promoting or inhibiting growth cone extension.. These developmental instructions establish appropriate RGC axon pathfinding, synaptogenesis, and retinotopographic mapping, and many of these cues are conserved across mammalian and nonmammalian vertebrate visual systems (Baier, 2013; Bollmann, 2019; Contreras et al., 2019; Erskine and Herrera, 2007; Harada et al., 2007; Seabrook et al., 2017; Siu and Murphy, 2018). However, the majority of efforts to date have focused on identifying these mechanisms in primary visual processing areas, and how RGCs target other critical brain regions remains incompletely understood (Barresi et al., 2010, 2005; Ross et al., 1992; Schnabl et al., 2021).

Zebrafish possess simplified visual circuitry that can be studied using tractable genetic tools. For example, zebrafish have monocular vision, and RGC axon connections occur only in the contralateral optic tectum (TeO) and pretectum (Pr) (Burrill and Easter, 1994; Robles et al., 2014). Due to the their neuroanatomical simplicity and speed of functional development, zebrafish were used in large-scale forward genetic screens to identify important genetic factors of visual system development (Baier et al., 1996; Bergeron et al., 2011; Brockerhoff et al., 1995; Karlstrom et al., 1996; Neuhauss et al., 1999; Seth et al., 2006; Trowe et al., 1996). These studies have furthered our understanding of conserved biological mechanisms underlying neurodevelopment and ocular function across vertebrates (Gestri et al., 2012; Sakai et al., 2018), yet like in many other systems, have largely focused on the primary center for visual processing, the TeO.

Most RGC axons terminate in the TeO, the neuroanatomical and functional equivalent structure of the mammalian superior colliculus (Sc). However, many of these RGC axons first terminate in the Pr before branching to send collaterals into the TeO (Burrill and Easter, 1994; Robles et al., 2014). Subclasses of RGCs have recently been defined based on their gene expression profiles (Kölsch et al., 2021; Nevin et al., 2010). Other studies have identified subclasses of RGC axons based on dendritic morphologies in the retina, position in the retina, their axon projection patterns and positions in the tectum (Förster et al., 2020; Robles et al., 2014), their post-synaptic partners (Gebhardt et al., 2013; Robles et al., 2013; Xiao et al., 2012, 2011), and their activity patterns to definitive features of visual stimuli (Kölsch et al., 2021; Zhou et al., 2020).

In zebrafish, RGC axons are further segregated into distinct retinorecipient neuropil regions called arborization fields (AFs) (Burrill and Easter, 1994; Robles et al., 2014), that are analogous in many ways to known visual processing regions in mammals (Baier and Wullimann, 2021; Yáñez et al., 2018). AFs are defined by their high density of presynaptic puncta and numbered 1 to 10 based on their proximity away from the midline optic chiasm (OC) and progression along the optic tract (Burrill and Easter, 1994; Robles et al., 2014). AFs 1-9 terminate in the Pr, and AF10 is in the TeO. Innate visual abilities have been differentially linked to distinct pretectal AFs, including computations underlying prey capture (Antinucci et al., 2019; Barker and Baier, 2015; Muto et al., 2017; Oldfield et al., 2020; Semmelhack et al., 2014; Wang et al., 2020, 2019), reflexive eye movements (Kubo et al., 2014; Naumann et al., 2016), optic flow (Kist and Portugues, 2019; Kramer et al., 2019; Temizer et al., 2015; Zhang et al., 2021), and motor asymmetry (Horstick et al., 2020). These behaviors are mediated by AFs 7, 9, 5, 6 and 3 respectively, while AFs, 1, 2, and 4 have no defined functions to date. While pretectal AF connectivity has been documented morphologically, and several pretectal AFs have unique roles in driving visually-mediated behaviors, there still remains a gap in knowledge of the cellular and molecular mechanisms that contribute to synaptogenesis of nine AFs across the Pr.

*Genomic screen homeobox 1 (Gsx1*) encodes a homeobox transcription factor that is expressed in the developing hypothalamus, olfactory bulb, cerebellum, hindbrain, spinal cord, TeO, and Pr (Bergeron et al., 2015; Cheesman and Eisen, 2004; Coltogirone et al., 2022; Satou et al., 2013). In mice, *gsx1* is expressed in the optic stalk at E11.5 (Valerius et al., 1995) and in precursor cells that give rise to the Sc at E13.5 (Bergeron et al., 2015). Integral roles for Gsx1 have been found in neural progenitor cell determination in ventral telencephalic regions in mice that ultimately give rise to cortical interneurons (Mizuguchi et al., 2006), and Gsx1 is known to limit proliferation and promote differentiation of glutamatergic and GABAergic interneurons in the mouse spinal cord (Pei et al., 2011). In zebrafish, *gsx1* promotes differentiation of both excitatory and inhibitory neurons in dorsal spinal cord progenitor domains (Satou et al., 2013). Examination of Gsx1 knockout (KO) mice have implicated Gsx1 in the hypothalamus and pituitary as contributing to normal hormone signaling that regulates body growth (Li et al., 1996). Similar roles in zebrafish have been identified in *gsx1* mutants (Coltogirone et al., 2022), however, zebrafish *gsx1* mutants are adult viable and fertile while Gsx1 KO mice do not survive long past 3 weeks of age (Li et al., 1996). Gsx1-positive neurons in the larval zebrafish brainstem modulate processing of acoustic stimuli in a paradigm related to endophenotypes associated with neurodevelopmental disorders (Bergeron et al., 2015; Tabor et al., 2018). While many roles for Gsx1 have been identified in early brain development and function across vertebrates, a role for Gsx1 in the developing visual system has not been explored despite its known expression there.

In this study we examine neuronal differentiation in the visual system in established *gsx1* zebrafish mutants (Coltogirone et al., 2022) and find that it is altered. We further characterize RGC axon connectivity patterns and identify visually mediated behavioral deficits in *gsx1* mutants that are consistent with developmental defects in the Pr. Lastly, we test cellular mechanisms contributing to RGC axon termination in the Pr using localized laser ablation. Our findings show for the first time that *gsx1* is required for termination of a subset of RGC axons and normal visual ability.

## RESULTS

### *gsx1* is required for glutamatergic neuron differentiation in the Pr but not neuron specification

Due to previously identified roles for *gsx1* in the differentiation of excitatory neurons in mouse and zebrafish (Pei et al., 2011; Satou et al., 2013), we first investigated glutamatergic neurons in the visual system using a transgenic line reporting expression of the vesicular glutamate transporter, *solute carrier family 17 member 6b (slc17a6b*, formerly *vglut2a* and *vglut2.1;* line *Tg(slc17a6b:DsRedf). slc17a6b* is a well-established marker for glutamatergic neurons in zebrafish (Kinkhabwala et al., 2011; Smear et al., 2007), with notable similarity to mammalian VGLUT2, belonging to a small group of proteins that mediate the uptake of glutamate by pre-synaptic vesicles (Fremeau et al., 2004, 2001; Takamori et al., 2001). Examination of *Tg(slc17a6b:DsRed*) at 6 dpf revealed a robust reduction in expression in *gsx1^y689^* within the anterolateral pretectal region compared to wildtypes (*gsx1^+/+^*), and heterozygotes (*gsx1^y689/+^*) (Fig. 1A-C).

**Fig. 1.**
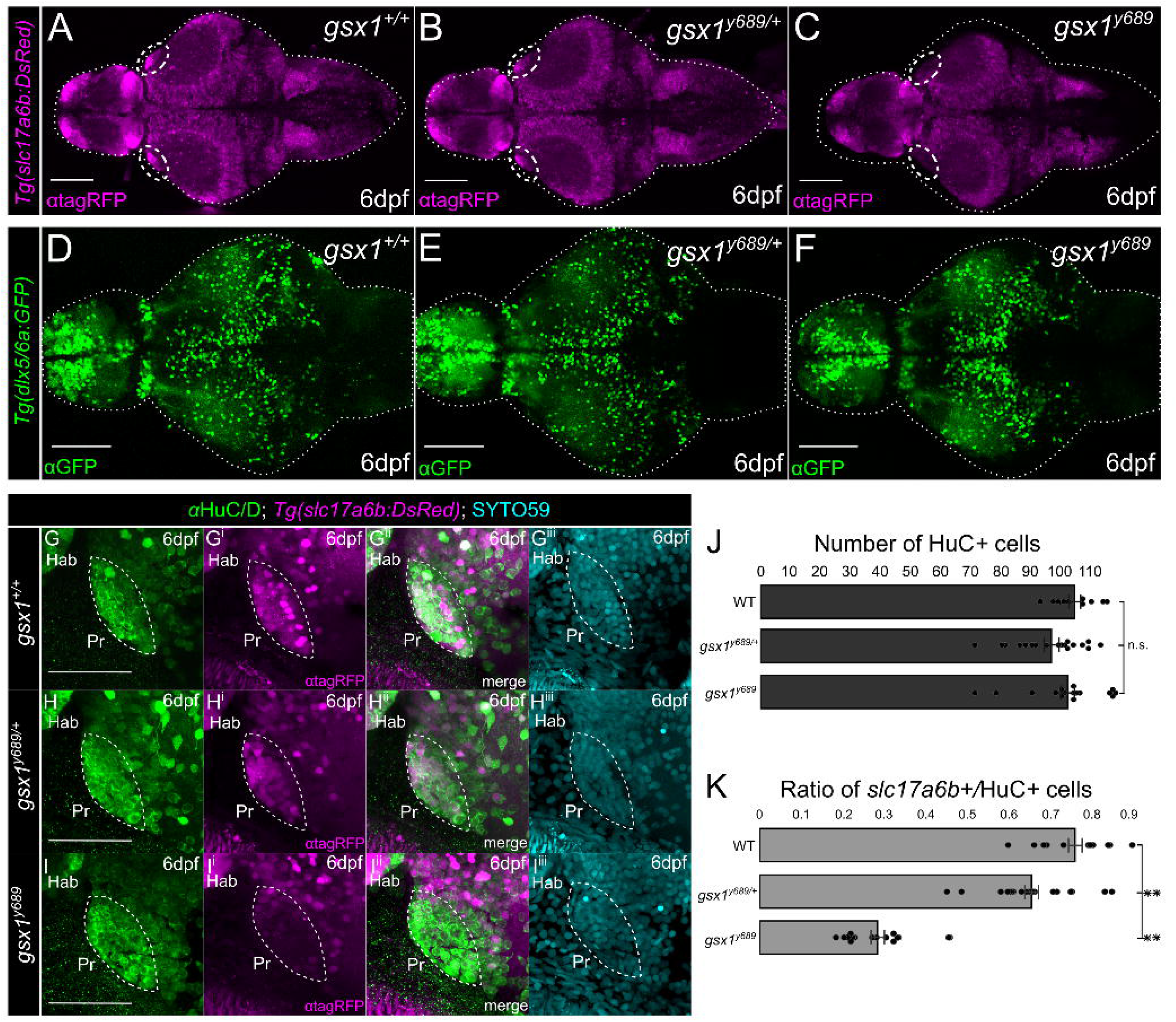
Examining excitatory and inhibitory neuron differentiation in *gsx1^y689^*. **(A-F)** Max projections of confocal z-stacks of *Tg(slc17a6b:DsRed*) and *Tg(dlx5a/6a:GFP*) at 6 dpf in *gsx1^+/+^* (wildtypes), *gsx1^y689/+^* (heterozygotes), and *gsx1^y689^* (mutants). Scalebar = 100μm. White dashed circle **(A-C)** outlines pretectal region missing *slc17a6b* in *gsx1^y689^*. **(G-I^iii^)** Max projection of confocal z-stacks (~20μm) of pretectal region using *Tg(slc17a6b:DsRed*) (magenta) and *HuC/D* (neurons, green) at 6 dpf. Cyan = SYTO59, nuclei. Scalebar = 50μm. **(J)** Bar graph with no significant differences found in total number of pretectal neurons by assessing *HuC/D* and SYTO59 staining across genotypes (single factor ANOVA F(2,43)=2.05, p=0.14). **(K)** Bar graph of average percentage of Pr *slc17a6b*-positive neuron counts out of *HuC/D* counts, *gsx1^+/+^* (n=11), *gsx1^y689/+^* (n=19), *gsx1^y689^* (n=16), F(2,43) = 3.21,*p*<.001. Post-hoc analysis of single factor ANOVA revealed *slc17a6b/HuC* ratio for *gsx1_+/+_* (79.72 ± 2.64) are significantly different from *gsx1^y689/+^* (63.95 ± 2.92,*p*<0.001) and *gsx1^y689^* (28.63±1.7,*p*<0.001). There is also a statistically significant difference between *slc17a6b/HuC* ratios *gsx1^y689/+^* and *gsx1^y689^* (*p*<0.001). Power factor greater than 80% for all genotypes, *p*<0.05.

We also assessed changes in inhibitory neurons in the visual system using *Tg(dlx5a/6a:GFP*), a transgenic line that reports expression of the transcription factor encoding genes that regulate inhibitory neuron differentiation in zebrafish (Hoffman et al., 2016; Yu et al., 2011) (Fig. 1D-F). We determined there was no obvious difference in expression of *dlx5a/6a* in the *gsx1^y689^* visual system; in fact, there were very few *dlx5a/6a* positive cells in the anterolateral Pr region in wildtypes. Closer examination of this Pr region in *gsx1^y689^* shows a loss of inhibitory neuron projections coming into the Pr and expressing *dlx5/6* (Fig. S1), thus we focused our efforts on quantifying *Tg(slc17a6b:DsRed*) expressing Pr neuron changes further.

We were able to assess if pretectal neurons were present in *gsx1^y689^* using an antibody that recognizes a protein expressed in neurons, HuC/D (Kim et al., 1996), and a fluorescent dye that binds DNA to demarcate nuclei, SYTO59 (Bradley et al., 2016). Quantification revealed that *gsx1* mutants have the same number of Pr neurons within the region devoid of *slc17a6b* expression (Fig. 1G-J), however, in *gsx1^y689^* only 28% of the neurons in that region express *slc17a6b*, while 66% of neurons were *slc17a6b*-positive in *gsx1*^y689/+^, and 76% of the total number of neurons were *slc17a6b*-positive in wildtypes (Fig. 1K). These results show that *gsx1* determines the neurochemical identity of a large proportion of pretectal neurons, but not pretectal neuron number. This is consistent with its role in mice to control the differentiation timing of neural precursors.

### Pretectal AF7 is predominantly disrupted in *gsx1* mutants

Retinorecipient cells within the Pr, thalamus, and TeO cluster based on their unique gene expression profiles that do not appear to change with loss of retinal input (Sherman et al., 2021). These findings and others suggest that genetic profiles are hard-wired in regions where RGC axons make connections and govern cellular mechanisms for the development and function of AFs (Baier, 2013; Brożko et al., 2022; Sherman et al., 2021). Due to identified changes in neuronal identity in the *gsx1* mutant Pr, we looked to next assess the morphology of incoming RGC axons using *Tg(atoh7:eGFP*), a transgenic line that demarcates all RGCs and their axons with GFP expression (Poggi et al., 2005). From our labeling and imaging, we determined RGC axons in *gsx1^y689^* fail to form AF7, AF8 may also be affected, and AF10 appears decreased in size, while other AFs including 3, 6, and 9 are still present at 6 dpf (Fig. 2A-C^ii^).

**Fig. 2.**
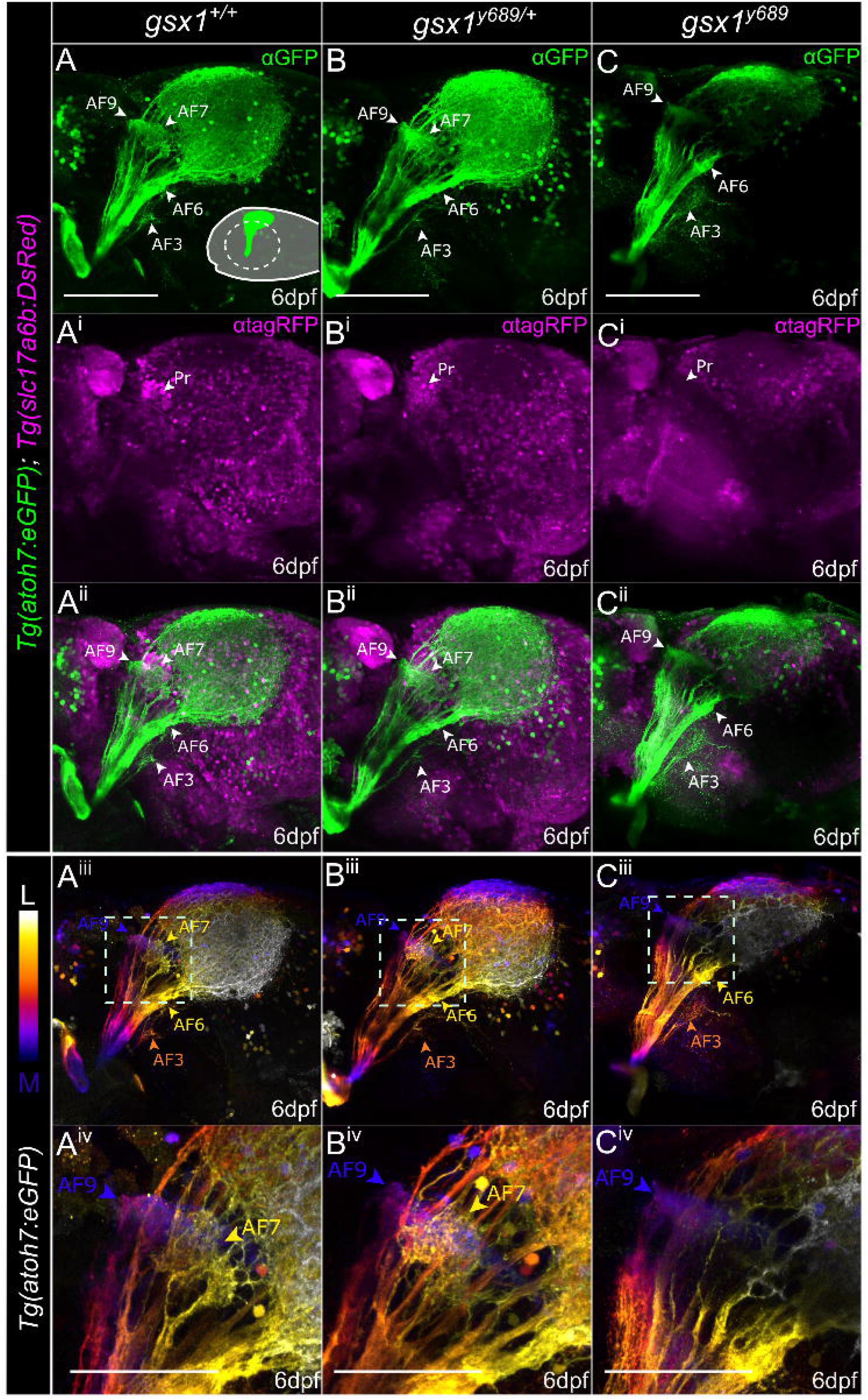
RGC axon termination is disrupted in *gsx1^y689^*. **(A-C)** Max projections of confocal z-stacks through the optic tectum from start of *Tg(atoh7:EGFP*) (green, RGC axons) labeling to end in **(A-A^ii^)** wildtypes (~85μm), **(B-B^ii^)** *gsx1^y689/+^* (~82μm), and **(C-C^ii^)** *gsx1* mutants (~80μm). *Tg(slc17a6b:DsRed*) labels glutamatergic neurons (magenta), and white arrowhead, Pr, points to pretectal region lacking *slc17a6b* expression in *gsx1^y689^*. Schematic in A displays lateral orientation with dashed circle indicating removed eye. Scalebar = 100μm. **(A^iii^-A^iv^, B^iii^-B^iv^, C^iii^-C^iv^)** Depth coded view of same samples in A-C^ii^ to provide reference for some AF regions with color coordinated arrowheads for depth, AF9 = dark blue, AF7 = yellow, AF3 = orange, AF6 = yellow. **(A^iv^, B^iv^, C^iv^)** Zoomed in view of the pretectal AFs from dashed outline box in A^iii^, B^iii^, C^iii^. Scalebar = 50μm. AF9 can be seen in blue across each genotype, while AF7 is absent in *gsx1^y689^*.

To better visualize individual AFs we used depth color coding of Pr RGC axons from deepest (AF9) to most superficial, medial to lateral (Fig. 2A^iii^-C^iv^). We further examined yellow colored RGC axons, at the level where AF7 is. In depth color-coded images for RGC axons, it appears AF9, which is known to develop near AF7, is unaffected across all genotypes demonstrating that this phenotype is specific to AFs in the AF7/8 region. AF7 and AF8 are too close in proximity to resolve by depth color coding. These findings show that with loss of *gsx1*, pretectal *slc17a6b* expression is reduced while neuron number stays the same, and RGC axons do not form terminals in the region of *slc17a6b* reduction near AF7/8 with additional defects present in RGC axon arbor size in AF10.

### RGC axon volumes and trajectories in *gsx1* mutants are only partly disrupted

Gsx1 KO mice and *gsx1* zebrafish mutants have decreased body size due to deficits in HPA signaling (Coltogirone et al., 2022; Li et al., 1996). Due to an observed reduction in AF10 in *gsx1^y689^* larval zebrafish at 6 dpf (Fig. 2), we were interested in assessing if the volume of RGC axon arbors could be measured to confirm a size decrease despite overall changes in mutant body size. We used *Tg(atoh7:eGFP*) to label all RGC axons and examine lateral views at 6 dpf in wildtype and *gsx1* mutants (Fig. 3A-B^i^). At 6 dpf, the optic tract extends from the midline to the contralateral side of the brain, and ventral AF1 and AF2 were not included in volume measurements. RGC axon neuropil regions from AF3 to AF10 in *gsx1^y689^* were found to be significantly smaller than wildtypes (Fig. 3C). As significant body size deficits in zebrafish *gsx1* mutants have not been identified until 14 dpf (Coltogirone et al., 2022), our findings support this axon phenotype at 6 dpf is likely irrespective of body size decrease in standard length.

**Fig. 3.**
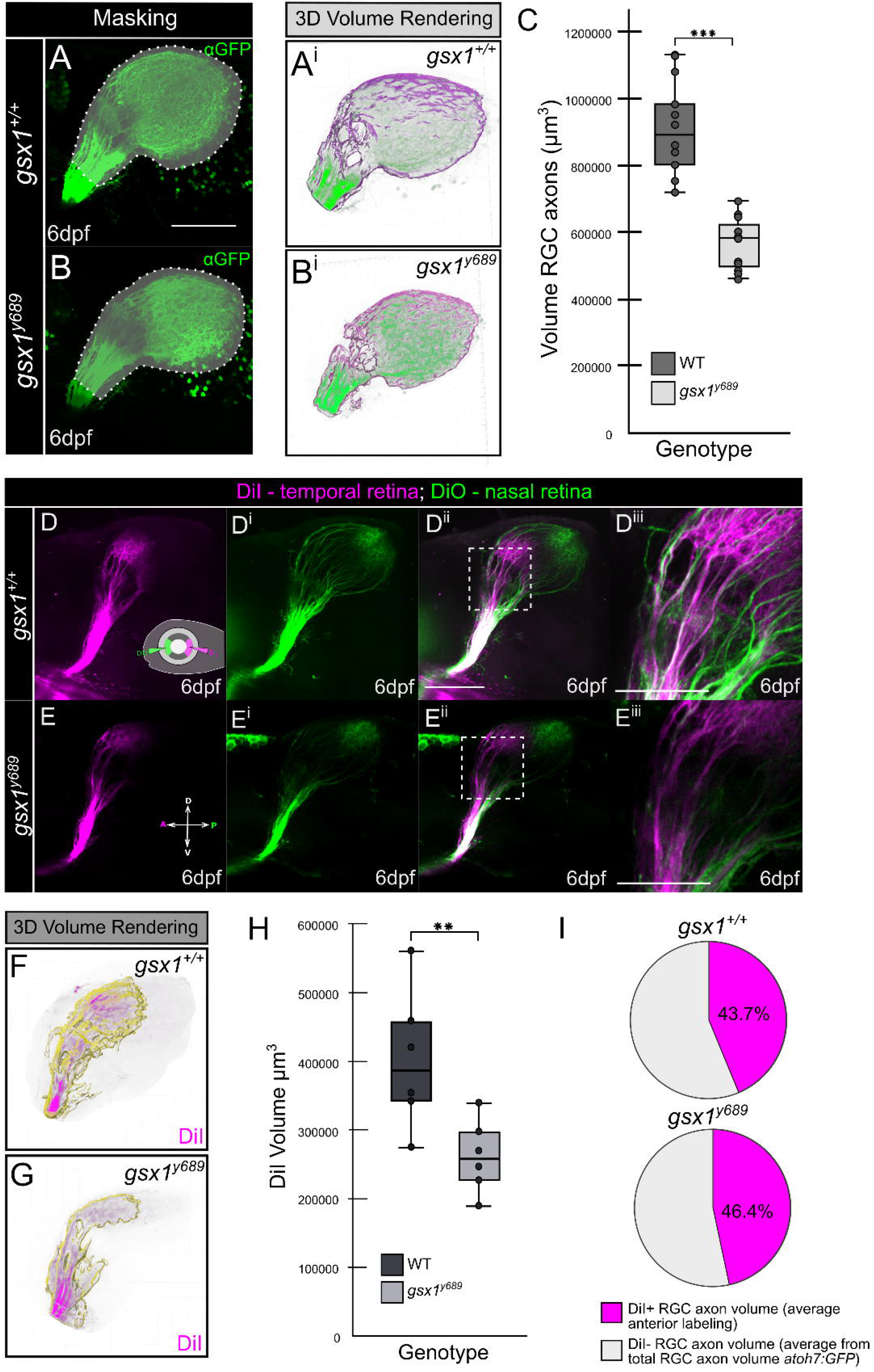
RGC axon volume examination in *gsx1* mutants. **(A)** Example max projection of a confocal z-stack in wildtypes and **(B)** *gsx1^y689^*, with *Tg(atoh7:eGFP*) labeling. Light grey trace around indicates region of interest and mask application. **(A^i^-B^i^)** Example 3D volume rendering taken from Imaris when Labkit is used. **(C)** Box and whisker plot with individual data points included for volumes. There was a significant difference in volume of full tectal lobe sizes between wildtype (M = 904100.00μm^3^, SD = 134984.32, SEM = 42685.79μm^3^, n = 10) and *gsx1* mutants (M = 564000.00μm^3^, SD = 80067.47, SEM = 24141.25μm^3^, n = 11); *t*(19) = 7.1,*p* < 0.001. Post-hoc power analysis = 100%. **(D-D^iii^)** Max projection of confocal z-stacks in wildtypes and **(E-E^iii^)** *gsx1^y689^*, with DiI (magenta) injected into the temporal retina and DiO (green) injected into the nasal retina. Scalebar = 100μm. **(D^iii^-E^iii^)** Zoomed in pretectal region with consistent loss of AF7 in *gsx1^y689^*. Scalebar = 50μm. **(F-G)** Same samples as D and E visualized through 3D volume rendering in Imaris software when Labkit is used. Yellow outline is volume, magenta is DiI labeling with mask to capture region of interest. **(H)** Box and whisker plot with individual data points included for volumes in WT and *gsx1^y689^*. There was a significant difference in volume of DiI labeling (magenta) between wildtype (M = 400833.33μm^3^, SD = 100882.94, SEM = 41185.29μm^3^, n = 6) and *gsx1* mutants (M = 261000.00μm^3^, SD = 53062.23, SEM = 21662.56μm^3^, n = 6); *t*(10) = 3.01, *p* < 0.01. Post-hoc power analysis = 100%. **(I)** Pie chart of RGC axon volume average *Tg(atoh7:GFP*) (taken from C) used to calculate a ratio of anterior DiI labeling from H, in wildtype (*gsx1*^+/+^) and *gsx1^y689^*.

Retinotopographic mapping in the zebrafish visual system has been classically examined by injection of lipophilic dyes into the retina in order to label specific subsets of anterior and posterior terminating RGC axons (Baier et al., 1996; Poulain and Chien, 2013; Robles et al., 2014). We anticipate that due to the loss of AF7, RGC axons in *gsx1^y689^* may terminate in off-target locations. To address this, DiI was injected into the temporal retina to distinctly label AF7 neuropil and anterior RGC axons in the Pr and TeO (Fig. 3D-E^iii^). Consistent with prior results, deficits in AF7 formation were detected in *gsx1^y689^* and quantification revealed that *gsx1* mutants had significantly smaller anterior RGC axon volumes (Fig. 3F-H). We next calculated a ratio of anterior DiI labeling out of the average RGC axon volume of *atoh7:eGFP* in AF3-10 in *gsx1^y689^* or wildtypes (Fig. 3I), in order to assess if ratios of anterior to posterior labeling were changed. Ratios indicated anterior DiI labeling occupied about 44% of total RGC axon volumes in wildtypes and about 46% in *gsx1^y689^*, revealing that there is a loss of anterior RGC axons in *gsx1^y689^* overall (Figure 3H) and that they maintain their general A/P retinotopographic termination zones despite deficits in the formation of AF7/8 specifically.

### Decreased prey capture in *gsx1* mutants

Functional analysis of AF7 has revealed its unequivocal role in performing prey capture (Antinucci et al., 2019; Barker and Baier, 2015; Kramer et al., 2019; Muto et al., 2017, 2013; Oldfield et al., 2020; Semmelhack et al., 2014). We aimed to functionally validate the observed defects in AF7 in *gsx1^y689^* by performing a prey capture assay in mutants compared to their wildtype cousins. We first determined an optimal amount of live rotifers to the number of larvae per testing dish. To do this, we measured the number of rotifers per milliliter (mL) before adding larvae (pre) and after 2 hours of exposure (post) (Fig. 4A-A^iii^). We then performed *in vitro* fertilization to obtain *gsx1* mutants and their wildtype cousins to assess prey capture at 7 dpf. *gsx1^y689^* had higher post counts of rotifers within their environment compared to wildtype cousins (Fig. 4B), indicating an overall reduction in prey capture success. These results functionally confirm our neuroanatomical findings that AF7 is absent in *gsx1^y689^*.

**Fig. 4.**
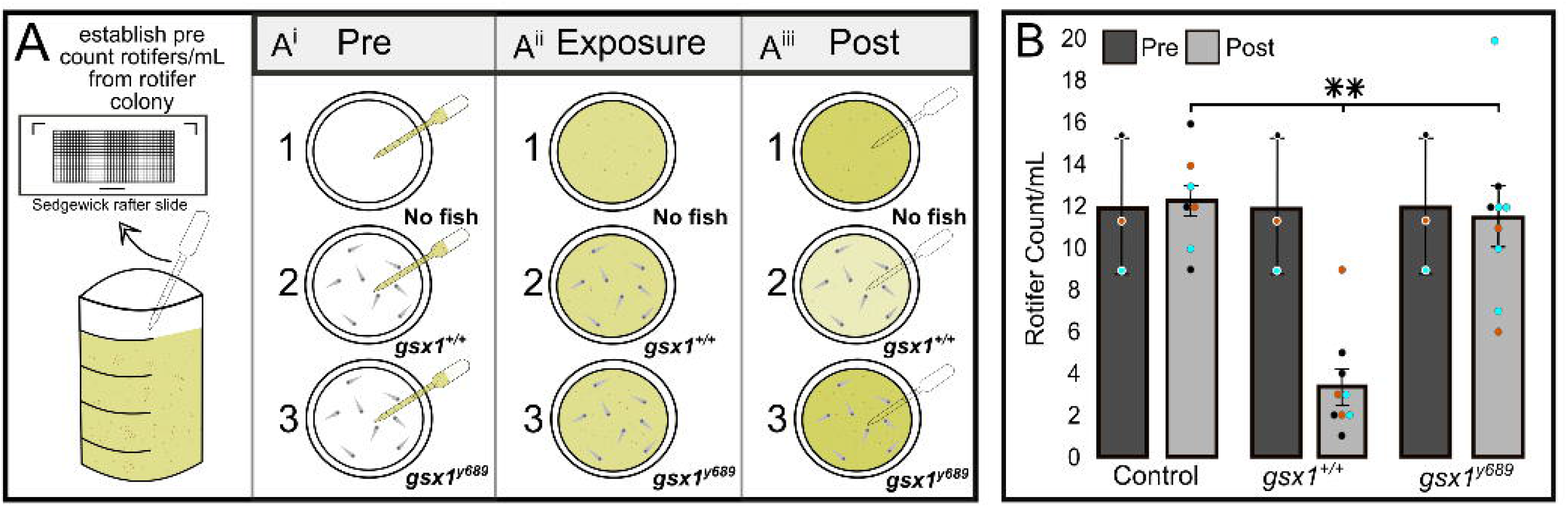
*gsx1* mutants have disrupted prey capture at 7dpf. **(A-A^iii^)** Experimental setup for prey capture assay; 8 7 dpf larvae per 6 cm dish, except in controls (no larvae). **(B)** Bar graph depicts rotifer/mL average across each group with individual data points showing all trials color coded (cyan, black, and orange) (n=3) for controls (n=7 dishes), wildtypes (n=9 dishes), and *gsx1^y689^* (n=9 dishes); pre (dark grey bars), post (light grey bars). Power analysis revealed 100% for each condition, *p*< 0.05. *gsx1^+/+^* post count of rotifers/mL were statistically different from controls and *gsx1^y689^*, (F(2,24)=[3.44], *p*<0.001). Error bars displayed as ±SEM. No significant differences found between control and *gsx1^y689^* post count analysis, *p*=0.63.

### *slc17a6b*-positive neurons are necessary for the formation of AF7

RGC axons follow complex sets of molecular cues upon eye exit and as they establish synapses on the contralateral side of the brain in the Pr and TeO (Baier, 2013). Therefore, we aimed to determine if the deficit to AF7 in *gsx1^y689^* is at the level of RGC axon termination or if additional disruptions exist before RGC axons reach the Pr. We found that *slc17a6b*-positive cells directly surround AF7 in wildtypes using *Tg(slc17a6b:DsRed);Tg(atoh7:eGFP*) in combination with anti-Synaptotagmin (Znp1) antibody, a marker for presynaptic terminals (Fig. S2). We next confirmed the results of our previous gene expression studies (Coltogirone et al., 2022) that *gsx1* is not expressed in the eye by performing RT-PCR on dissected eyes only or heads without eyes (Fig. S3A-A^ii^). We also examined the morphology of the retinal ganglion cell layer (GCL) at 6 dpf using *Tg(isl2b:GFP*) to label RGCs (Fig. S3B-D^ii^). RGCs within the eye appear qualitatively and quantitatively normal across *gsx1* genotypes (Fig. S3E).

*gsx1* is known to be present in the TeO and Pr by 48 hpf and even earlier in neural precursor regions in the forebrain (Coltogirone et al., 2022), consistent with the developmental period when RGC axons cross the forebrain midline, forming the optic chiasm (Barresi et al., 2005). We further examined optic chiasm crossing and optic nerve formation at 48 hpf and no changes were detected across all genotypes (Fig. S4). These results align with previous findings that *gsx1^y689^* does not have changes in early (1-2 dpf) forebrain patterning gene expression such as *dlx2a* and *dlx2b*, thus the cell and molecular substrate that RGC axons grow out upon is largely normal (Coltogirone et al., 2022). We hypothesize that the loss of AF7 is specifically related in some way to the loss of *slc17a6b-expression* in *gsx1^y689^* pretectal neurons.

In order to assess if *slc17a6b*-positive cells in the Pr are involved in RGC axon termination and formation of AF7, we performed unilateral removal of pretectal *slc17a6b*-positive neurons at 72 hpf using 2-photon laser-ablation in *Tg(atoh7:eGFP);Tg(slc17a6b:DsRed);gsx1^+/+^* and examined the establishment of RGC axon terminals at 7 dpf (Fig. 5). The zebrafish visual system allows manipulations to be confined to one pretectal lobe, leaving the ipsilateral side intact and is ideal to be used for phenotypic comparison. At 72 hpf, the visual system is still developing, however, presumptive RGC axon neuropil regions can be seen, and the pretectal region of interest is morphologically distinct and identifiable using *Tg(slc17a6b:DsRed*) (Fig. 5C). RGC axon patterns at 7 dpf show that AF7 formation was disrupted after pretectal *slc17a6b*-positive neuron ablation compared to non-ablated controls and the intact Pr side (Fig. 5I-K). The phenotypes for 7 dpf AF7 disruptions are specifically based on depth color coding (Fig. 5J^iv^,K^iv^), and images indicate other pretectal AFs are unaffected by *slc17a6b*-positive neuron ablation.

**Fig. 5.**
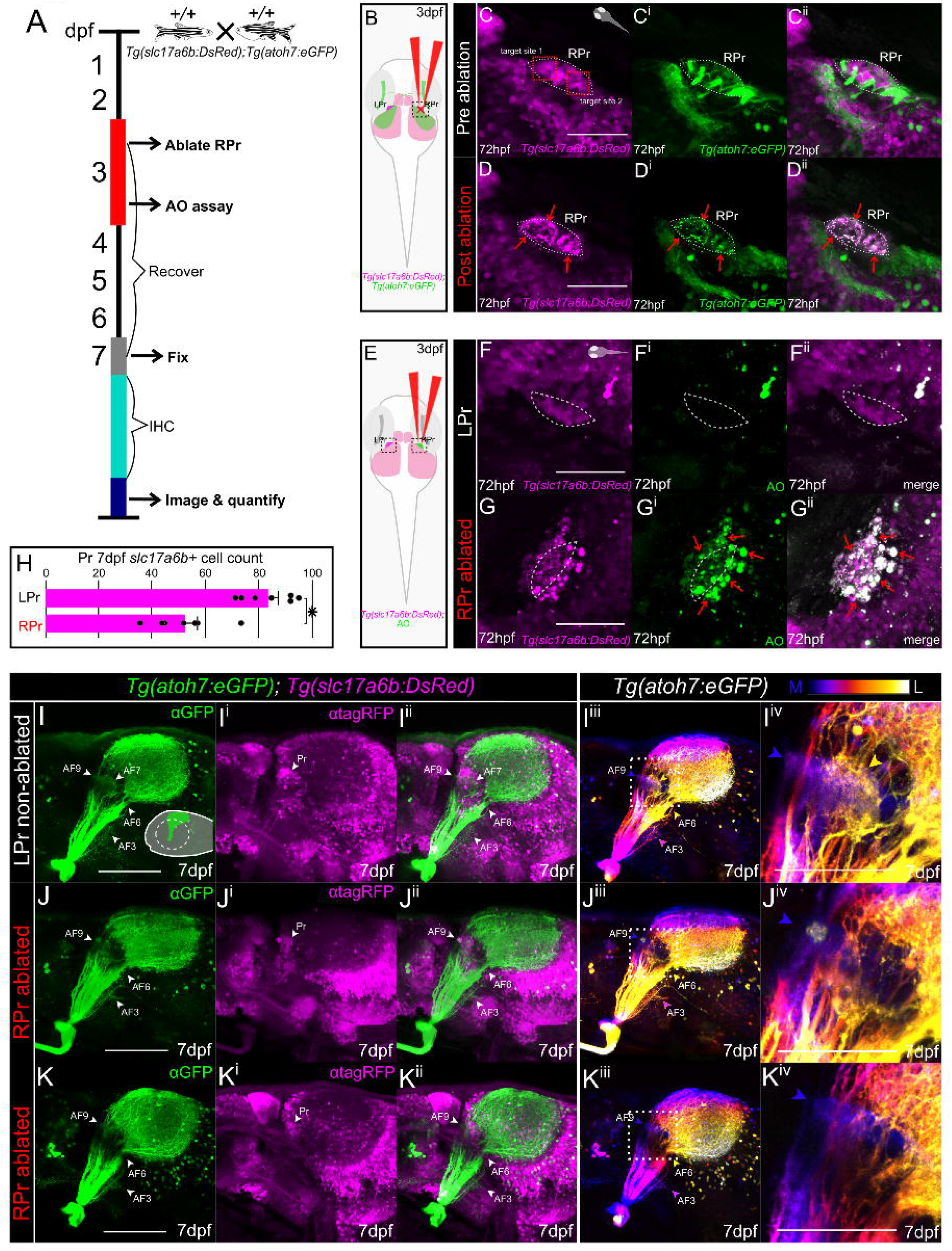
*slc17a6b-positive* pretectal neurons are required for AF7 formation. **(A)** Experimental timeline: *gsx1^+/+^ Tg(slc17a6b:DsRed);Tg(atoh7:eGFP*) were raised until 72hpf when they undergo unilateral ablation of the right pretectal (RPr) region. An acridine orange (AO) assay is performed on some samples at 72hpf, hours post-ablation. Other ablated samples recover until 7 dpf. At 7 dpf samples are fixed and further undergo immunohistochemistry (IHC). **(B)** Schematic of unilateral RPr ablation in *gsx1^+/+^ Tg(slc17a6b:DsRed);Tg(atoh7:eGFP*) (magenta and green, respectively) at 72 hpf. LPr = left pretectum. **(C-C^ii^)** Pre-ablation of 72 hpf max projection of 2P z-stack through RPr separated by individual channels of *Tg(atoh7:eGFP*), green, *Tg(slc17a6b:DsRed*), magenta, and merged image channels. Two ablation sites are targeted and shown in C, red boxes. Dashed white outline indicates pretectal region. Orientation of sample is in right top corner. Scalebar = 50μm. **(D-D^ii^)** Post RPr ablation of same sample at 72 hpf, red arrows indicate displacement of the fluorescent proteins. **(E)** Schematic of imaging planes in black dashed boxes after unilateral RPr ablation and then following AO staining. **(F-G)** AO staining compared to intact LPr in the same RPr ablated sample, using *Tg(slc17a6b:DsRed*) (magenta) while red arrowheads indicate increased acridine orange (green) in ablated side. **(H)** Bar graph of quantified *slc17a6b-positive* neurons at 7 dpf in the LPr and RPr following 72 hpf RPr ablation. The ablated RPr showed statistically significant decreases in *slc17a6b-positive* neurons compared to LPr side (n=7), *t*(13)=0.96, *p*<0.001. **(I-K)** Lateral 7 dpf max projections of confocal z-stacks of RGC axons in *Tg(atoh7:eGFP*) and *Tg(slc17a6b:DsRed*) and the same images depth color coded from medial (blue) to lateral (white), providing AF visualization, blue = AF9, yellow = AF7. Scalebar = 100μm. **(I-I^iv^)** Control (no ablation, n=6), orientation schematic in right bottom corner. **(J-K^iv^)** 72hpf unilateral RPr ablated resulting in AF7 disruption at 7 dpf (n=11/11). **(I^iv^, J^iv^, K^iv^)** Zoomed in image from adjacent images with white dashed boxes for visualization of RGC axon AF patterns. Scalebar = 50μm.

We confirmed our *slc17a6b*-positive neuron ablation technique through acridine orange (AO) staining on the ablated side of the Pr compared to the non-ablated side at 72 hpf, roughly 4 hours after ablation (Fig. 5F-G^ii^). Pretectal cell death on the ablated side was also seen at 6 dpf (3 days post ablation), however, AO staining is undetectable at 7 dpf (4 days post ablation), indicating that between 6 and 7dpf, the Pr cells may undergo cycling, consistent with findings from other studies examining these time points (Kita et al., 2015b). To understand how pretectal *slc17a6b*-positive neurons were impacted at 7 dpf, we quantified them on both the ablated and non-ablated Pr sides within the same samples, revealing a significant reduction in *slc17a6b*-positive neurons on the ablated pretectal side (Fig. 5H). While pretectal *slc17a6b* reduction post ablation is statistically significant, loss of *slc17a6b*-positive neurons does not completely match the reduction seen in *gsx1* mutants at 6 dpf (Fig. 1H). However, it was sufficient to prevent AF7 from forming and is one full day after our initial quantification in *gsx1^y689^*. We conclude that pretectal *slc17a6b-expressing* neurons are required during development for RGC axons to terminate within the neuropil region for AF7.

### RGC axons can establish proper pretectal retinotopographic positions following the ablation of *slc17a6b*-positive neurons

Since RGC axons are near pretectal cells by 72 hpf, the possibility that they are also partly ablated upon *slc17a6b*-positive neuron ablation arises. However, studies have shown that the optic nerve can re-establish proper connections following transection by 4 days post injury (dpi) without additional RGC proliferation or cell death (Harvey et al., 2019). This finding establishes that RGC axons likely have the capability over the course of four days to find their proper position within the Pr if ablated. To examine this with our laser-ablation technique, we ablated a region of the optic nerve near AF6 using *Tg(atoh7:eGFP*) and examined RGC axon patterns 4 days post-ablation (Fig. S5). RGC axon patterns for AF6 are normal in ablated larvae at 7 dpf in comparison to controls (Fig. S5D-E). These findings further support that pretectal *slc17a6b* expressing cells are involved in RGC axon termination and formation of AF7, and ablated RGC axons at 72 hpf by our laser-focused methods have the capacity to reform connections over the course of four days.

To further investigate termination errors seen within our ablation experiments, we assessed retinotectal pathfinding following unilateral ablation of the pretectal cells. Ventro-temporal RGCs within the eye are known to project to AF7 and branch to form additional terminals within the anterior TeO (Robles et al., 2014). In order to assess if RGC axons occupy anatomically correct retinotectal termination zones following right-sided pretectal (RPr) cellular ablation using *Tg(HuC:GFP*) at 72 hpf, a lipophilic dye (DiI) was injected into the temporal region of the controlateral retina at 7 dpf to evaluate the formation of AF7 and the anterior Pr/TeO (Fig. 6A). Following ablation to the RPr, DiI-tracing via the left eye confirmed all RGC axons cross the optic chiasm and terminate within the anterior portion of the RPr/TeO consistent with control experiments at 7 dpf (Fig. 6B-C). We also observed that RGC axons do not show off-target projections within the Pr/TeO (Fig. 6D-E). RGC axons within RPr ablated samples do not appear to have termination within posterior RTeO positions or additional locations where RGC axon projections can be visualized. Zoomed in images with depth color code for the RPr/TeO also indicate a phenocopy of *gsx1^y689^*, specific to AF7 following unilateral RPr ablations (Fig. 6D^iii^,E^iii^), as seen in previous experiments. Our ablation techniques using *Tg(HuC:GFP*) instead of *Tg(slc17a6b:DsRed*) produce similar phenotypes. These experiments show that RGC axons re-establish proper retinotopographic and retinotectal locations following the ablation of pretectal cells and only AF7 is affected.

**Fig. 6.**
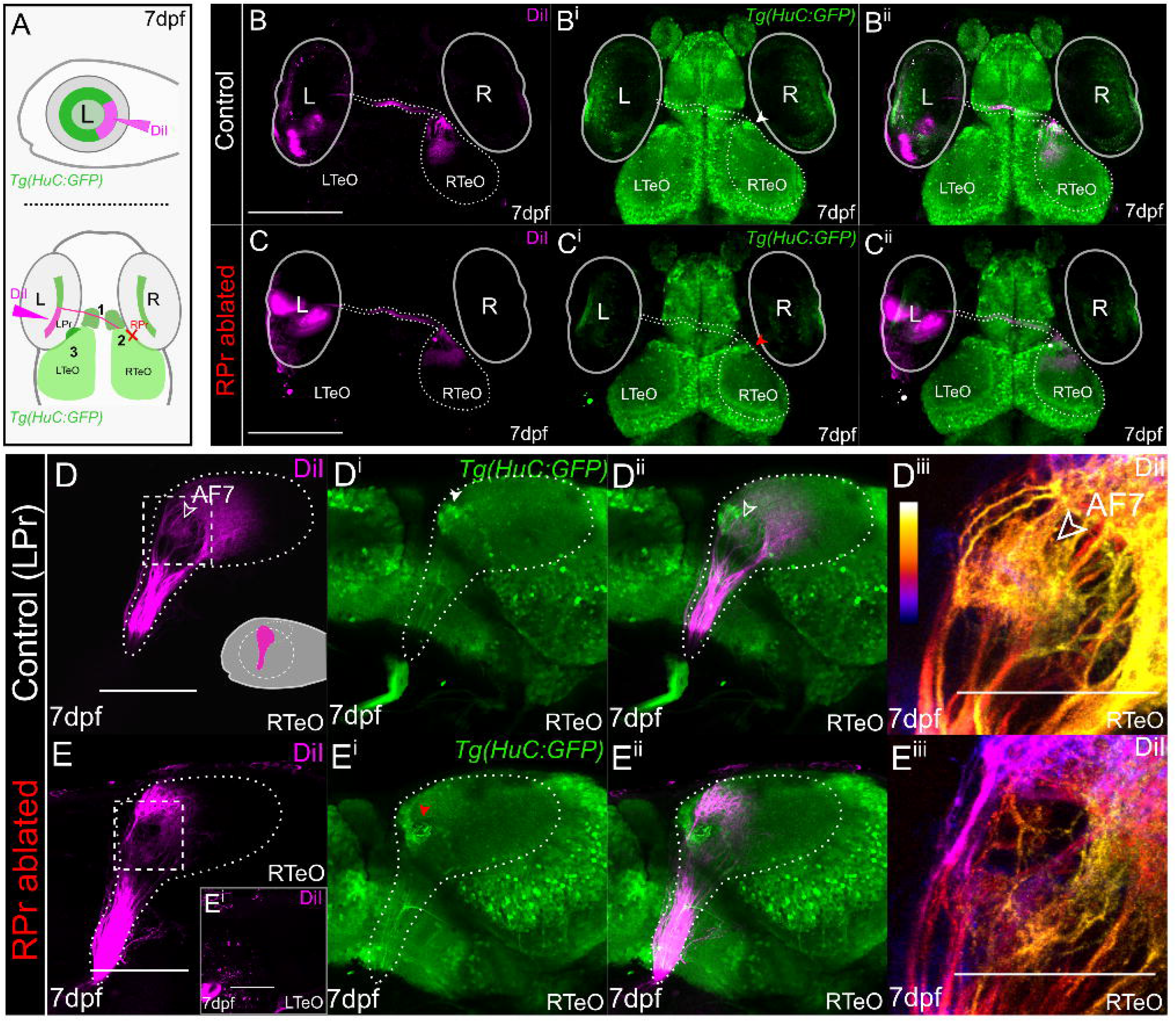
Retinotopographic order is confirmed via lipophilic dye labeling after unilateral pretectal ablation. **(A)** Schematics indicating orientation and location of left eye (L) DiI injections and areas of RGC axon examination to the contralateral right TeO/Pr. Three black numbers indicate regions to examine labeling after ablation; 1) optic chiasm crossing, 2) anterior tectal labeling on contralateral lobe from injection, and 3) potential DiI labeled RGC axons within ipsilateral lobe from injected eye (left optic tectum = LTeO). R = right eye. **(B-B^ii^)** Max projections of 2P z-stacks (~200μm) in control non-ablated samples at 7dpf (n=10). DiI = magenta, green = *Tg(HuC:GFP*). **(C-C^ii^)** Unilateral RPr ablations in *Tg(HuC:GFP*) wildtypes at 72 hpf, fixed at 7 dpf and DiI injected in temporal retina. Max projections of 2P z-stacks show proper optic chiasm crossing, anterior RGC axon arborization, and no RGC axons are seen targeting LTeO or elsewhere on injected side (n=11). Red arrowhead indicates RPr that was ablated at 72hpf. Scalebar = 200μm. **(D-D^iii^)** Max projection of confocal z-stacks of RTeO in lateral orientation showing DiI labeling in control (LPr) non-ablated samples (n=5), (~85μm). Schematic of left eye DiI injection. White arrowhead indicates RPr region and red arrowhead indicates ablated RPr region. Scalebar = 100μm. **(E-E^iii^)** Lateral orientation with left eye removed of max projected confocal z-stacks across RPr ablated samples and DiI labeling in RTeO (~85μm). **(D^iii^, E^iii^)** DiI image from D and E in white dashed box is depth color coded from medial (blue) to lateral (white) and zoomed in showing loss of AF7 (yellow). Scalebar = 50μm.

## DISCUSSION

In this study, *gsx1* is shown for the first time to contribute to the differentiation of glutamatergic neurons in the visual system that aide in establishing proper eye to brain neural circuit connectivity during development. We found that in *gsx1* mutants, RGC axon termination is disrupted including the formation of AF7. In wildtype larvae, *slc17a6b*-expressing neurons directly surround AF7, and we establish through cellular ablation and RGC axon tracing techniques that the formation of AF7 is dependent on pretectal *slc17a6b-positive* neurons. We also confirm that *gsx1* is not expressed in the eye, and *gsx1* mutants have normal GCL morphology in the retina. We further examine RGC axon pathfinding in the forebrain and identify that *gsx1^y689^* have typical patterns of RGC axon exit from the eye and optic chiasm formation. Following the removal of pretectal *slc17a6b-positive* neurons during early development, we establish that RGC retinotectal order is maintained at larval stages, and deficiencies are further narrowed to RGC axon termination and synaptogenesis errors in AF7. Axon branching from this region may also be affected given the reduction we observed in AF10 size in the TeO. Additionally, we reveal that the visually mediated behavior of prey capture is decreased with the loss of *gsx1*, which is consistent with the functional profiles of AF7 and its absence in these mutants. In summary, pretectal neurons expressing *slc17a6b* are dependent on *gsx1* for differentiation, and these *slc17a6b-positive* neurons influence RGC axon termination at AF7 to drive visual behavior (Fig. 7). Recent data based on functional and neuroanatomical findings suggests a great majority of the neurons that provide local (postsynaptic) communication to AF7, reside within the migrated pretectal region (M1) or superficial parvocellular pretectal (PSp) nucleus (Baier and Wullimann, 2021; Butler et al., 1991; Yáñez et al., 2018). All together we demonstrate the importance of *gsx1* for the development and function of visual neural circuits for the first time in any vertebrate.

**Fig. 7.**
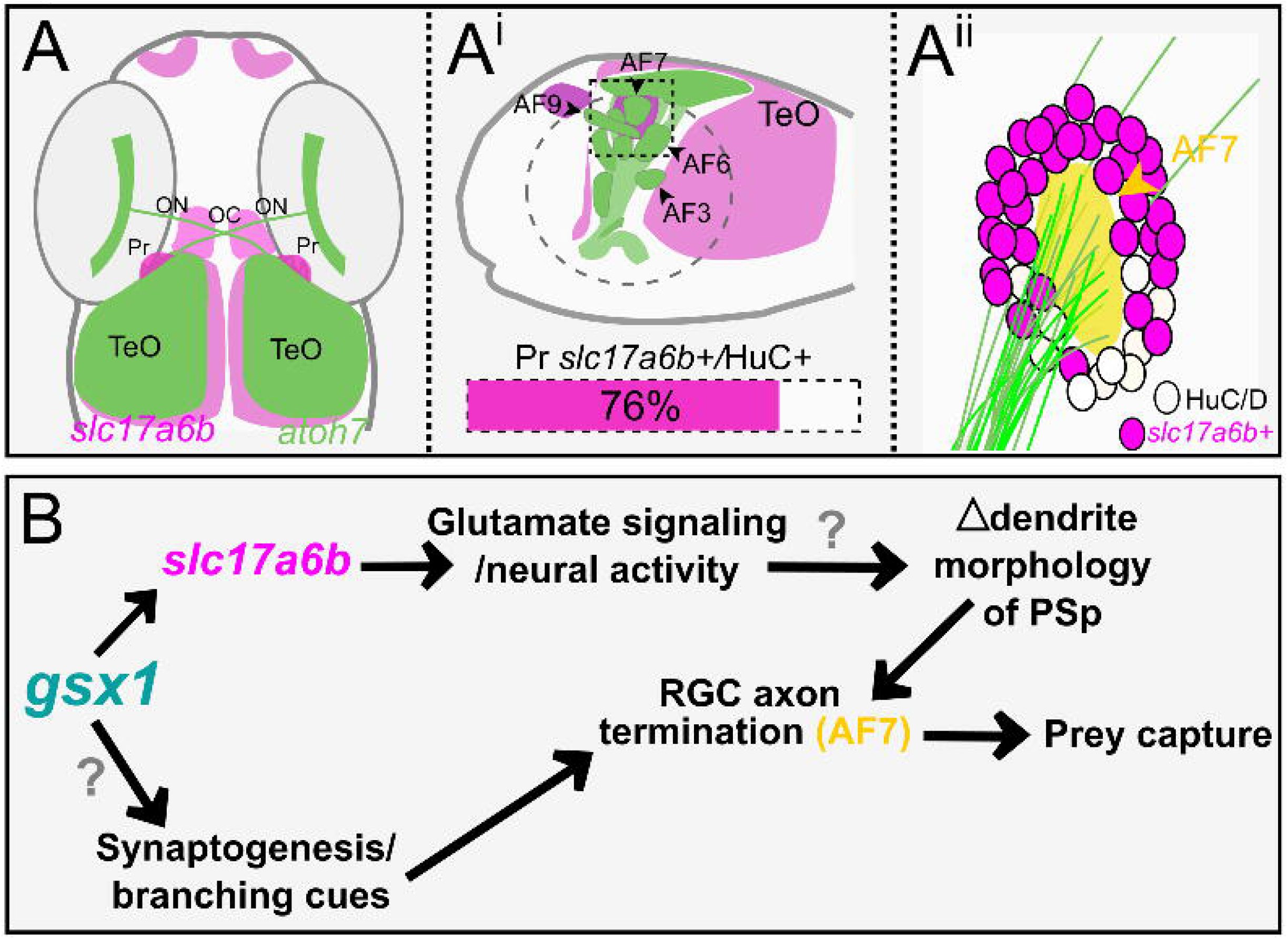
A model for *gsx1* function in the visual system. **(A)** Summary schematic of *slc17a6b* expression (magenta) and RGC axon patterning via *atoh7* expression (green) at 6 dpf in wildtypes (*gsx1^+/+^*). **(A^i^)** Lateral view of the TeO and pretectal arborization fields (AFs) (adapted from (Robles et al., 2014)) with *slc17a6b* expression (magenta). Wildtype pretectal *slc17a6b/HuC/D+* cell quantity is shown with magenta bar. **(A^ii^)** Zoomed in schematic of RGC axons for AF7 surrounded by *slc17a6b-positive* cells from the outlined box in Ai. **(B)** Model of how *gsx1* contributes to pretectal neuronal identity and neural circuit assembly. Parvocellular superficial pretectal nucleus (PSp).

### Pretectal *slc17a6b*-expressing neurons are involved in the development of AF7

Our findings are in line with identified roles for Gsx1 in the differentiation of neuronal subtypes in mouse and zebrafish (Mizuguchi et al., 2006; Pei et al., 2011; Satou et al., 2013). Recent findings in mice indicate that the reintroduction of a lentivirus encoding Gsx1 can mediate functional recovery of neurons following spinal cord injury by increasing the number of glutamatergic and cholinergic neurons (Patel et al., 2021). Our results indicate that with the loss of *gsx1*, mutants have changes to specific regions targeted by RGC axons. RGC axon termination errors are specific to AF7/8 and AF10, suggesting that the molecular identity of these recipient neurons is key to shaping incoming RGC axon synapses. Although *slc17a6b* is widely used to mark the majority of glutamatergic neurons in zebrafish, other vesicular glutamate transporters do exist within zebrafish, including *slc17a6a* (formally *vglut2b), slc17a7a* (formally *vglut1*) and *slc17a8* (formally *vglut3*) (Smear et al., 2007). However, *slc17a6a* is not uniquely detectable by antibody or transgenic line thus far and is reported to have overlapping expression profiles with that of *slc17a6b. slc17a7a* is also weakly expressed within the visual system, and *slc17a8* has been reported to have minimal expression, and therefore they were not included in our investigation. Despite its broad expression, the defects we observed in *slc17a6b* were very specific in only the regions where *gsx1* is also expressed and beyond the Pr extended into the brainstem and hypothalamus.

Global loss of *slc17a6b* in the *blumenkohl (blu*) mutant results in reduced visual ability (Smear et al., 2007). In addition, expanded RGC axon arbors with enlarged receptive fields are observed in the *blu* TeO (Smear et al., 2007). This study confirmed that other glutamate transporters were still available but were not enough to overcome visual fatigue and expansion of RGC axon arbors with loss of *slc17a6b*, highlighting glutamatergic neurotransmission at the retinotectal synapse is mainly modulated through *slc17a6b*. Previous studies have also investigated retinotectal glutamate signaling in zebrafish through bath application of antagonists for the receptors N-methyl-D-aspartate (NMDA), α-amino-3-hydroxy-5-methyl-4-isoxazolepropionic acid (AMPA), and kainate (KA) (Schmidt et al., 2000; Smear et al., 2007), with resulting phenotypes consistent with *blu* and enlarged RGC axon arbors if blockage occurred during different critical periods of visual system development. However, these experiments have examined RGC axon patterns following global changes to glutamate signaling, namely on both pre and postsynaptic sides of the terminal synapses. Our findings show that *slc17a6b* is decreased selectively in the Pr in *gsx1^y689^* while the eye remains normal due to the absence of *gsx1* expression there. Often, RGC axon branches have been shown to be rapidly eliminated when postsynaptic connections are not found (Kita et al., 2015b; Kutsarova et al., 2017), which may explain loss of AF7. Our model is the first to implicate *slc17a6b* in the Pr alone as having a role in the development of specific AFs, requiring glutamatergic neurotransmission on the postsynaptic side of pretectal neural circuits to provide incoming RGC axons on the presynaptic side with cues for termination. Neurons that are less active during development may have less suitable dendrite morphology for normal synaptogenesis to occur, leading us to test this and other hypotheses about these *slc17a6b*-positive pretectal cells surrounding AF7 further as summarized in Figure 7B. It is also possible that transcriptional target genes of Gsx1 include known and novel axon termination, synaptogenesis, and/or branching cues for RGC axons in the Pr that are worth further investigation in light of this subtle but impactful phenotype.

### Visual neuropil region size is altered, but RGC axon topography is maintained

We discovered that the total RGC axon volume in *gsx1* mutants was reduced and can be easily visualized by quantifying three-dimensional axon volume from AF3 to AF10. Pretectal AFs terminate in anterior retinotectal regions and can be labeled with lipophilic dyes injected into the temporal retina (Robles et al., 2014). We hypothesized that *slc17a6b* expressing pretectal neurons may produce stop signals for incoming RGC axons to terminate and upon loss of these signals, that these axons may occupy and target alternative regions. We analyzed retinotectal targeting and discovered that RGC axons do not expand across anterior locations despite AF7 deficits, nor do they extend more into posterior TeO regions. While our results show that anterior labeling is decreased but stays in normal proportion compared to the full RGC axon volume, our techniques might not be sensitive enough to determine subtle changes in RGC axon targeting across the retinorecipient field of the TeO and Pr. Alternatively, AF7 designated axons may occupy unconventional anterior regions such as another pretectal AF, which would not be fully accounted for in this analysis. These subtle defects may be more easily detected using newly developed transgenic tools (Kölsch et al., 2021) to label where AF7 axons ultimately reside in *gsx1^y689^*. It would be interesting to explore time points earlier in development with more advanced genetic and imaging tools in hand to pinpoint when RGC axon volume differences are initiated in *gsx1^y689^* and if critical windows shape morphological patterns during axon targeting and refinement of connections in the absence of *gsx1*. It is also important to note that the AF7 neuropil region appears to receive a lot of inhibitory input from *dlx5/6*-positive neurons, and these connections also fail to form in our *gsx1* mutants, adding to the important roles worth dissecting out for *gsx1* in the formation of the visual system.

### Visually mediated dysfunction linked to neurodevelopmental changes

Prey capture requires neural circuits for visual perception, motor control, and decision making, which together give rise to the innate visually mediated response (Muto and Kawakami, 2013). The existence and activity of AF7 have been highly linked to prey capture abilities, and this region within our studies was shown to be morphologically impacted with loss of *gsx1*. Both the left and right pretectal regions may function together during different phases of prey capture, and integration of interhemispheric tectal neuron populations serves to mediate motor coordination programs (Bianco and Engert, 2015; Gebhardt et al., 2019; Nikolaou and Meyer, 2015; Oldfield et al., 2020). We suspect that there are other changes to be identified in neuronal differentiation in the TeO based on the documented expression of *gsx1* localized to the outer edges of the tectal neuropil (Coltogirone et al., 2022), which likely further contribute to positional changes to incoming RGC axons based on the reduction of AF10 that we measured. AF10 reduction could also be due to axon branching failures upstream in the Pr region affected in *gsx1* mutants. These connections may play a role in sensorimotor processing and integration. As *gsx1* is also expressed in the hypothalamus which is another main driving center in the brain for feeding behavior, we must also consider the role that it plays there. However, these and other changes throughout the CNS related to loss of *gsx1* are difficult to dissect out given the initial deficit that we identified in AF7 without more advanced genetic tools.

Since *gsx1^y689^* are adult viable, they make an excellent model to assess how early neurodevelopmental changes in the visual system influence later life behaviors that rely on an intact visual ability such as social behaviors and learning. We might also assess how disruptions in select AFs influence the function of others nearby. For example, AF9 has been tied to neural circuits underlying eye tracking in zebrafish via functional imaging in combination with optokinetic response (OKR) analysis (Muto et al., 2005; Robles et al., 2014). Since AF9 appears morphologically normal in our studies, we predict normal OKR in *gsx1^y689^*, however, based on close proximity to AF7 during development, there could be cross talk between these two regions that would be absent in our mutants. In addition, studies have shown that zebrafish without *gsx1*-expressing neurons and mice Gsx1 KOs have disrupted prepulse inhibition (PPI) (Bergeron et al., 2015; Tabor et al., 2018), a sensory gating behavior often disrupted in neurodevelopmental disorders (NDDs) such as schizophrenia (Braff et al., 1978). Clinical populations of patients diagnosed with schizophrenia also have deficits related to the inhibition of saccadic eye movement reflexes, called anti-saccade errors (Lencer et al., 2017). Further exploration of a role for *gsx1* in the development of visually mediated behaviors such as examination of saccadic eye movements may elucidate additional functions for *gsx1* in sensory processing that are widely linked to NDDs (Lencer et al., 2017; Levy et al., 2010).

Upon functional loss of *gsx1*, there are likely changes to known and novel molecular cues in the Pr and TeO influencing RGC axon termination, synaptogenesis, and branching. Our ablation techniques for assessing pretectal *slc17a6b*-positive cells early in development disrupt these cues that guide RGC axons to their proper retinotopographic locations in a very localized way. Future studies to understand the full repertoire of molecular networks that *gsx1* regulates will uncover potentially conserved developmental mechanisms that influence RGC axon guidance and termination within visual neural circuits. Importantly, these studies link *gsx1* to sensory processing disruptions in the visual system in addition to the ones it has already been linked to in the auditory system. Across the CNS, *gsx1* appears to be a key player in sensory processing neural circuit development and function.

## MATERIALS AND METHODS

### Zebrafish Husbandry

All aspects of this study were approved by the West Virginia University IACUC. Adult zebrafish were maintained as described previously (Coltogirone et al., 2022) at 25-28°C on a 14h/10h light/dark cycle. All embryos were staged under a dissecting scope (Kimmel, 1993; Kimmel et al., 1995) and raised in E3 embryo media (pH 7.4; 0.005M NaCl, 0.00017M KCl, 0.00033M CaCl, 0.00033M MgSO4.7H_2_0, 1.5 mM HEPES) at 28.5°C in an incubator with a 14h/10h light/dark cycle. Embryos and larvae used for histochemical techniques had E3 exchanged for 0.003% phenylthiourea (PTU) in E3 after 6 hpf to prevent pigmentation. Wildtype (WT, *gsx1^+/+^*), *gsx1* heterozygotes (*gsx1^y689/+^*), and *gsx1* mutant (*gsx1^y689/y689^*) fish lines in a TL background (Coltogirone et al., 2022), and transgenic *Tg(slc17a6b:DsRed*) (Kinkhabwala et al., 2011)*, Tg(atoh7:eGFP*) (Poggi et al., 2005), *Tg(HuC:GFP*) (Park et al., 2000), and *Tg(isl2b:GFP*) were used for these studies. For non-transgenic and non-mutant-based experiments, TL (Tupfel long fin) were used.

### Immunohistochemistry (IHC)

For histochemical techniques, *gsx1^y689/+^* in various transgenic backgrounds were in-crossed to obtain offspring. Desired ages and expression patterns of transgenes are obtained before fixation in 4% PFA in 1x phosphate buffered saline (PBS) at 4°C overnight. The following day, embryos and larvae are washed in 1x PBS, and undergo genotyping from dissected tail tissue, and brain and/or eye dissection is performed as needed before continuing with immunolabeling in order to achieve better penetration of antibodies. Repeating 1x phosphate TritonX (PTx) washes are completed at room temperature and then followed by 100% cold acetone treatment at −20C. Samples were then incubated in a blocking solution (5% normal goat serum, 1% DMSO in 1x PTx) for 1 hour at room temperature and then provided fresh block and incubation with primary antibodies (Table 1) at 4°C at a range of 2-4 overnights. Primary antibodies are then removed, and samples are washed with 1x PTx. Secondary antibodies are added (Table 1) with the same concentration of blocking reagents used previously, overnight at 4°C.

**Table 1.**
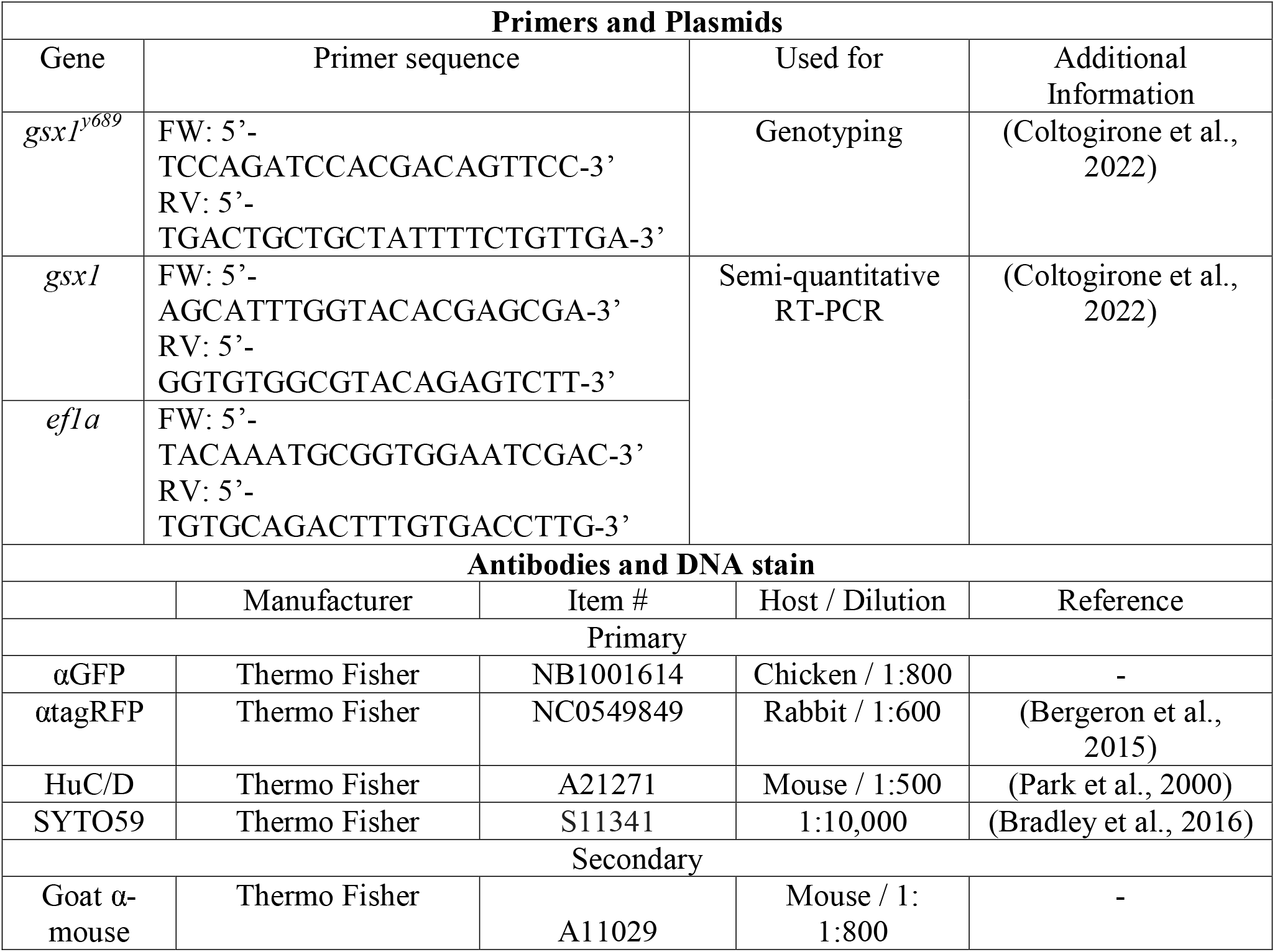
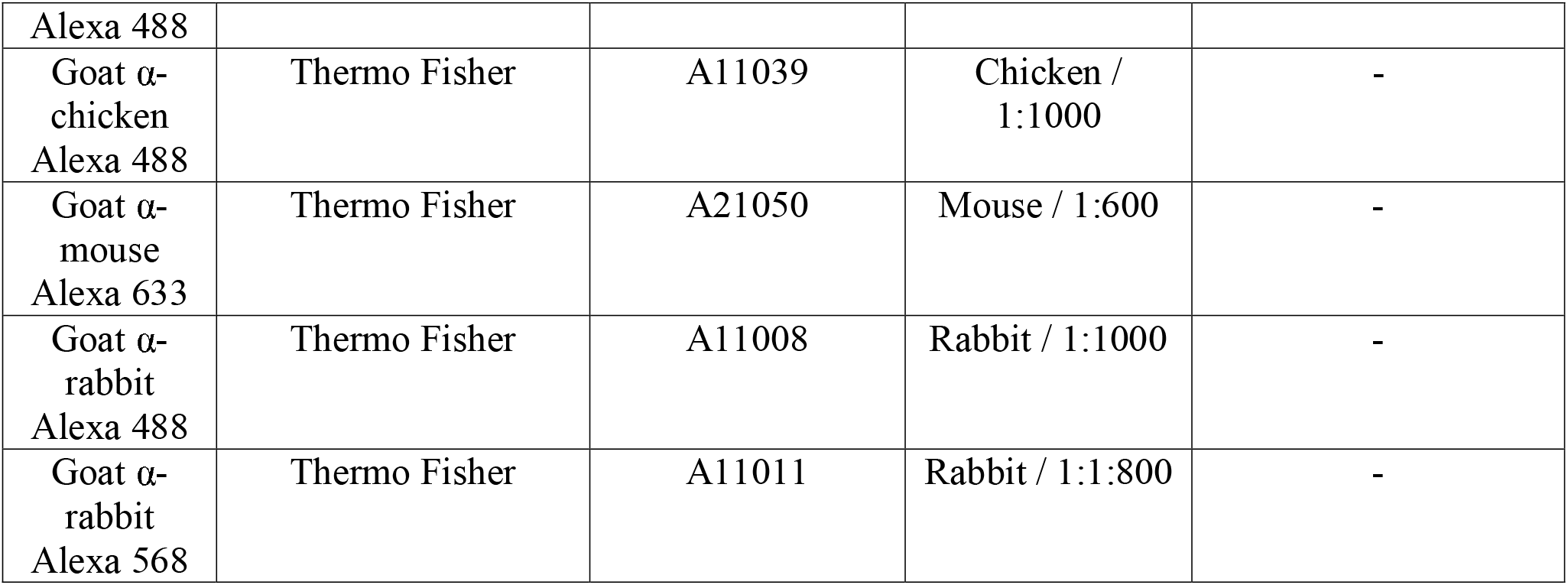
List of reagents used.

### Prey Capture Assay

Adult *gsx1^+/+^* or their siblings, *gsx1^y689^*, underwent separate *in vitro* fertilization (IVF) rounds to acquire two groups of cousin larvae raised side-by-side for behavior testing. 12mL of E3 was exchanged daily (n=30 individuals per 6 cm dish) until 7 dpf. At 7 dpf, a 100 mL sample of the rotifer colony was acquired, filtered, and diluted in E3. A Sedgewick Rafter counting slide was used to assess how many rotifers were present per mL, establishing a pre-count that was recorded and added to individual dishes. Groups of n=8 larvae were then transferred to each individual dish. One dish contained no larvae to account for multiplying rotifers. Dishes were placed into our light and temperature-controlled fish room. After 2 hours, a 1 mL sample is taken from each dish and counted for a post-count of rotifers per mL. Three replicates of individual IVF rounds with the same experimental setup were obtained. Pre and post counts of rotifers per mL were analyzed using a Students T-test, p=0.05, to assess prey capture success following the 2 hours of exposure. Post counts of each condition of rotifers per mL were taken blind to the experimenter and power analysis of α = 0.05 and 80% (ClinCalc) was sufficient for each group of statistical testing.

### Genotyping

Genotyping followed the protocol in Coltogirone et al., 2022 for *gsx1^y689^*. A small tail portion from the most posterior end was dissected from adults, larvae, or embryos to use in DNA prep. Tails were denatured at 95°C in DNA lysis buffer (10mM Tris pH 7.5, 50mM KCl, 0.3% Tween20, 0.3% Triton X, 1mM EDTA) and digested at 55°C for at least 2h with 10mg/mL proteinase-K (Omega) (1:5 volume dilution). The proteinase-K was heat-deactivated at 95°C before the DNA sample was used in a standard DreamTaq (Thermo Fisher) PCR reaction with gene-specific primers for *gsx1* (Table 1). Amplified fragments were analyzed using 4% agarose gel electrophoresis; *gsx1^+/+^* (wildtype) appear as one larger band (140bp), mutants appear as one smaller band (129bp), and heterozygotes appear as two bands, one at each size.

### RT-PCR for *gsx1* mRNA

Embryos and larvae obtained from TL crosses were raised to the desired ages (30 hpf, 48 hpf, 6 dpf) and stored in RNAlater (Sigma, #MKCL5657). Total RNA was extracted from 10 dissected eyes and 10 dissected heads without eyes at each age using a phenol chloroform extraction method with TRI-Reagent (Invitrogen). 2μg of cDNA prepared using a 2-step RT-PCR kit (Invitrogen) was used in 28 cycles of PCR with PlatinumTaq (Invitrogen) and intron-spanning gene-specific primers for *gsx1* and *ef1α* (Table 1). Fragments were visualized and imaged using a FluorChemQ imager (ProteinSimple) on a 3% agarose gel with SYBR Safe DNA gel stain (Invitrogen) using a blue light transilluminator (Clare Chemical Research). A Nanodrop fluorospectrometer was used to assess RNA quality before RT-PCR and gel electrophoresis.

### Lipophilic Dye Injections

7 dpf *gsx1^+/+^ Tg(slc17a6b:DsRed*) or *Tg(HuC:GFP*) larvae were fixed overnight in 4% PFA following 4 days of recovery from unilateral pretectal ablations at 72 hpf with fresh 0.003% PTU in E3 exchanged daily. Larvae were laterally mounted in 1.5% agarose in 1x PBS with the eye contralateral to the ablated side directed surface up. DiI (1mg/mL dissolved in ethanol) (Invitrogen, cat. #D3911) was in injected with a WPI Nanoliter microinjector into temporal-ventral locations within the retina. Samples were incubated for 1 hour at 37°C and then left to diffuse overnight at 4°C. Samples were removed from agarose and mounted dorsally and laterally in 1X PBS. Samples were imaged on a 2-photon or confocal microscope within 24 hours of DiI injection to visualize retinotopographic patterns.

For retinotectal mapping, *gsx1^y689/+^* adult zebrafish were in-crossed and offspring raised to 6 dpf and fixed with 4% PFA overnight. Larvae were laterally mounted in 1.5% agarose in 1x PBS. DiO (1mg/mL dissolved in DMF) (Invitrogen, cat. # D275) was injected into the nasal retina and samples were left in a 37°C incubator for 1 hour. The same samples were then injected with DiI (1mg/mL dissolved in ethanol) (Invitrogen, cat. #D3911) into temporal-ventral locations within the retina and then left to diffuse overnight at 4°C. Samples were removed from agarose and mounted laterally in 1X PBS and scanned at 2μm depths through the full DiI and DiO labeling on the confocal microscope within 24 hours of injection.

### Cryostat Retinal Sections and Quantification

6 dpf larvae (*Tg(isl2b:GFP);gsx1^y689/+^* in-cross) were fixed in 4% PFA overnight at 4°C. Tissue was prepped by sinking larvae in a 25% sucrose in 1x PBS solution, then 35% sucrose in 1X PBS and mounted in Optical Cutting Temperature (OCT Clear, Fisher HealthCare, 4585). Sections were taken on a Leica CM1850 Cryostat at 12μm through the entire retina based on published protocols (Uribe and Gross, 2007). Staining procedures for sections followed (Uribe and Gross, 2007), and anti-GFP antibody and SYTO59 staining were used (Table 1). Images were taken on a laser scanning confocal microscope and analyzed using Vaa3D software to count cells.

### Imaging: Confocal, 2-photon, and Epifluorescent Microscopy

Images were taken on an Olympus BX61 confocal microscope with Fluoview FV1000 software with imaging objectives 20X, 40X, 60X oil immersion, or 40X silicon oil immersion. Scientifica VivoScope 2-photon microscope with Spectra Physics Mai Tai HP tunable lasers and ScanImage software was used to perform 2-photon laser-mediated ablation experiments, *in vivo* imaging of acridine orange staining, and DiI injections after photo-ablation. Zeiss Axiozoom V16 Microscope with fluorescence was used to sort transgene expression and examine antibody labeling following IHC.

### Quantification of TeO neuropil and DiI RGC axon labeling volumes

Lateral confocal images using *Tg(atoh7:eGFP*) (RGC axons) and SYTO59 (nuclear label) were acquired by imaging through the entire tectal lobe from start of RGC axon fluorescence to end. The same imaging volume was taken following DiI and DiO lipophilic dye injections in *gsx1^y689/+^* in-crossed fish. Images were taken at 2μm per slice and stored as.oib files. Images were converted using Imaris 9.3 file converter and input into Imaris 9.9 (http://www.bitplane.com/imaris/imaris, Oxford Instruments). Masks were taken of region of interest starting at the base after AF1 and AF2 labeling to eliminate optic nerve measurements but include the Pr (AF3-9) and TeO (AF10). A mask of RGC axon labeling was then input into Labkit pixel classifier (ImageJ FIJI) and volume rendering was accomplished via a machine learning algorithm. Computed results were sent back to Imaris to obtain a reading of volume for our region of interest. Imaris volume rendering and capture was taken blind to the researcher by sample genotype. Volume data was analyzed in IBM SPSS (2021).

### Unilateral Pretectal Axon and Cell Ablations

For cell ablation, wildtype (*gsx1^+/+^*) adult fish with transgenes *Tg(slc17a6b:DsRed);Tg(atoh7:eGFP*) were in-crossed and sorted for both GFP and RFP expression at 48 hpf. PTU was added at 6 hpf to prevent pigmentation and embryos were raised in an incubator at 28.5□C. At 72 hpf (3 dpf) zebrafish were anesthetized with MS-222 and mounted in 1% low-melt agarose in E3 in a 4-well custom glass slide chamber. Each embryo was located using epifluorescence with a 16x objective (Nikon) and then located on the 2-photon at 920nm with 5x magnification to visualization left or right pretectal regions pre-ablation. Scans were taken pre-ablation with PMTs (~450nm) detecting GFP/RFP in *Tg(slc17a6b:DsRed*) and *Tg(atoh7:eGFP*) at ~15-18μW laser power. Target locations in unilateral pretectal regions were zoomed into using digital zoom features to 65x and laser power was increased to ~94-99μW, exposing the region of *slc17a6b-positive* cells for 1 second. Additional target regions were identified by zooming out and turning down laser power to locate the regions and then zooming back into 65x to expose the second pretectal region of *slc17a6b-positive* cells in the same way. Post-ablation images were taken at 5x to assess post-ablation success. Displacement of the fluorescent protein can be visualized in post-ablation images to identify successful targeting, and acridine orange was included to visualize dying cells. Larva were taken out of agarose carefully and recovered in E3 until 0.003% PTU is returned a few hours later. Experiments to assess larvae health after unilateral ablations determined that following ablations, 100% of swim bladders inflated and overall health is maintained through 7 dpf. Ablated larval health was compared to controls that are also mounted and unmounted in 1% agarose in E3 as a sham ablation.

For axon ablation control experiments, wildtype adult zebrafish carrying *Tg(atoh7:eGFP*) were in-crossed and embryos were collected to be placed in 0.003% PTU in E3 with daily exchanges of this rearing solution. At 72 hpf, the right eye of each larva was enucleated and they were placed immediately in Hanks’ solution (Thermo Fisher) to recover for 2 hours. This allowed for targeting of RGC axon regions on the unilateral side. After recovery, enucleated larvae were mounted in a customer chamber in 1% agarose in E3 with the enucleated side facing the glass immersion slide. Larvae were visualized with epifluorescence for precise mounting, and the ventral region of the optic nerve was targeted for ablation. Ablation parameters were the same for this experiment as used previously. Following ablation, larvae are un-mounted and placed back in 0.003% PTU in E3 to recover for 4 days with fresh media exchange and overall health checks performed daily. Control, enucleated larvae are treated the same way as experimental ablated samples except without ablation.

### Acridine Orange Assay

72 hpf *Tg(slc17a6b:DsRed*) wildtype zebrafish underwent unilateral pretectal photo-ablation through the aforementioned parameters and were left to recover in E3 for 2 hours. A cell death assay was used to assess photo-ablation success 2 hours post ablation with 5μg/mL of acridine orange (AO) (Ribeiro et al., 2007) (Sigma) added to each 1mL dish of E3. Ablated larvae (n = 6) and control non-ablated larvae (n = 6) were live mounted in 1% low melt agarose in E3 with 1mL MS-222 added following AO staining and re-imaged on the 2-photon microscope, zoomed into both right and left pretectal regions at a wavelength of 920nm. PMTs detected both GFP (AO) and RFP (*Tg(slc17a6b:DsRed)*) (~450nm). Cell death assays were also performed the same way at 6 dpf (3 days post ablation) and 7 dpf (4 days post ablation) to visualize if AO staining was still present after multiple days of recovery. Parameters for larval recovery after ablation and image acquisition were the same for each experiment.

### Image Processing, Cell Counting, and Statistical Analysis

Images were max projected unless indicated in Fiji ImageJ with brightness and contrast adjusted for image quality. Vaa3D (v.3.20) was used to count positive cell labeling in both pretectal sides using individual image captures zoomed into right and left pretectal lobes. Pretectal cell counts did not indicate differences in HuC/D or *slc17a6b* positive cells across right or left side and therefore were included grouped as all representative pretectal lobes (one counted lobe, n = 1). Optic nerve measurements were taken in Fiji ImageJ from max projected z-stack image files and averaged across 3 individual size measurements for both the right and left optic nerve at the location of the nerve leaving the retina. SPSS one-way ANOVAs and post hoc Tukey’s tests (α = 0.05) were used to determine significant differences between genotypes of pretectal cell counts (*slc17a6b, HuC/D*) and optic nerve measurements. Fiji ImageJ Plugin, Z-stack Depth ColorCode 0.0.2, was used to depth code for arborization field visualization and then max projected to produce z-stacks. Post-ablation depth coded images were analyzed for presence or absence of AF7 formation by examination of coded color for yellow and using blinded analysis of ablated or control lateral view max projection z-stacks. AF9 was matched in depth code to begin with blue coloring, indicating medial optic tecta location and the beginning of AF visualization throughout all samples. Student’s *t* tests were used for statistical comparison of *slc17a6b* counts in ablated versus non-ablated Pr on the contralateral sides in the same larvae. Outliers for statistical testing were determined by SPSS software for descriptive statistics and confirmed via GraphPad (Grubb’s test). Image analysis was done blind of the genotype. Power analysis of α = 0.05 and 80% (ClinCalc) was sufficient for each group of statistical testing.

## ACKNOWLEDGEMENTS

The authors wish to thank Eric Horstick, PhD at WVU for assistance with the 2-photon laser ablations, Imaris software, and review of this manuscript. We also thank Jeffrey Gross, PhD and his lab at UTexas, Austin for expert advice and assistance preparing and analyzing our retina cryosections and helpful discussions about this project. We would also like to thank Malia Miller at WVU for training on the 2-photon imaging system. Lastly, we thank Andrew Dacks, PhD at WVU for assistance and training on the confocal microscope and advice about immunohistochemistry and thoughtful review of our manuscript in preparation.

## CONTRIBUTIONS TO THE MANUSCRIPT

SAB and ARS conceived and performed the study, wrote and revised the manuscript, and analyzed and presented the data. RS gathered initial data on the Pr cell types and RGC axon morphology. RLP performed *in vitro* fertilization for prey capture studies and provided expert fish lines assistance for this research.

## COMPETING INTERESTS

No competing interests declared by any author.

## FUNDING

SAB funding that supported this work includes WVU Department of Biology lab startup funds, NICHD R15HD101974-01A1, and NIGMS P20GM144230-01. ARS was also supported by NIGMS-T32, 5 T32 GM 81741-10, WVU Eberly College of Arts and Sciences Doctoral Research Award, and the WVU Office of Academic Affairs Doctoral Research Award.

## SUPPLEMENTARY FIGURES

**SFig. 1.**
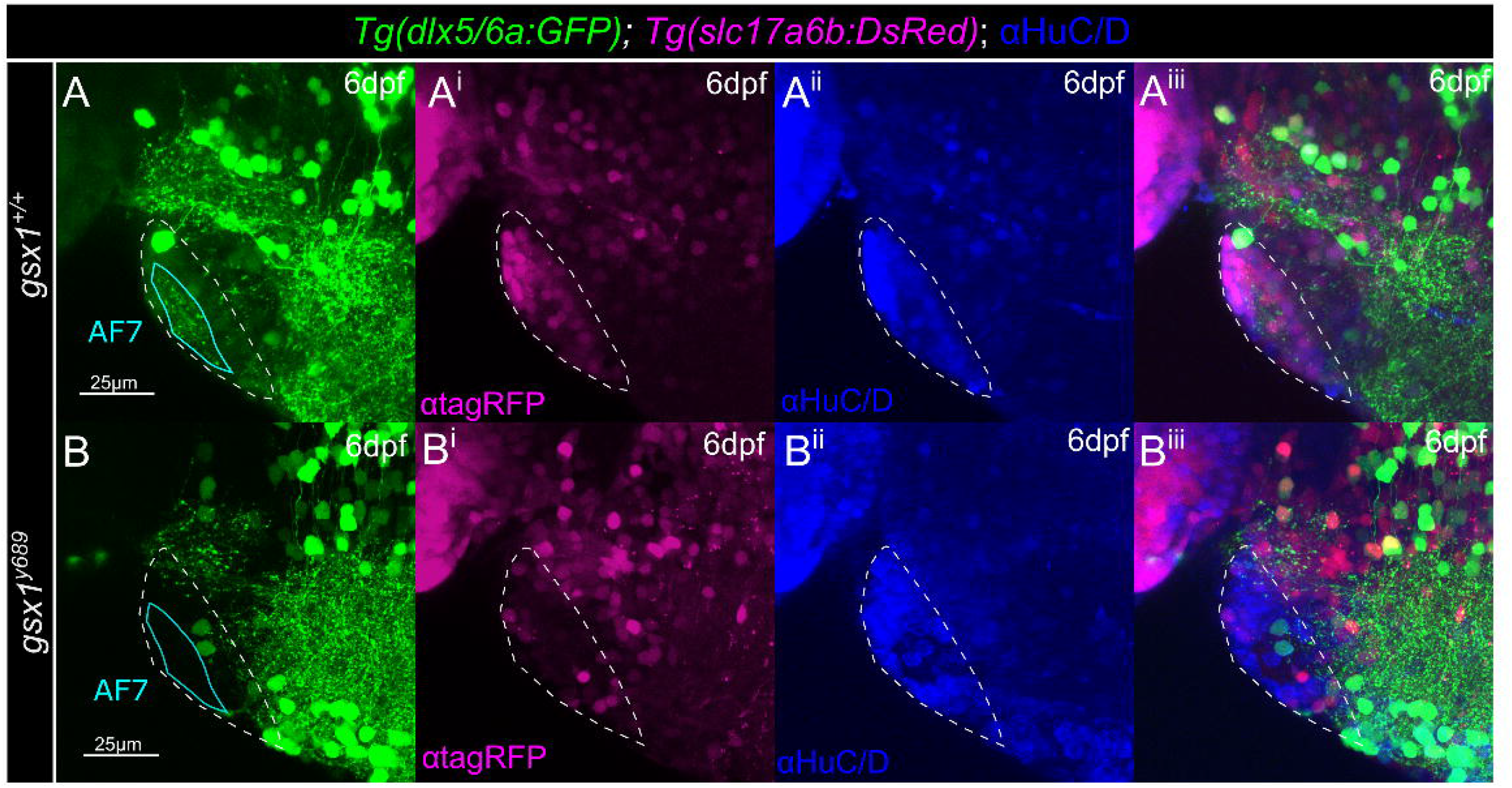
*Tg(dlx5/6a:GFP*) is a marker for inhibitory neuron connections to the pretectum that are missing in *gsx1^y689^*. **(A-B)** Max projection of confocal z-stacks through pretectal region (~20μm), in **(A-A^iii^)***gsx1^+/+^* and **(B-B^iii^)** *gsx1^y689^*. HuC/D = blue, *Tg(dlx5/6a:GFP*) = green, *Tg(slc17a6b:DsRed*) = magenta. White dashed line outlines pretectal region. Cyan line indicates arborization field (AF7) region with inhibitory connections to this region that appear to be missing in *gsx1* mutants compared to *gsx1^+/+^*.

**SFig. 2.**
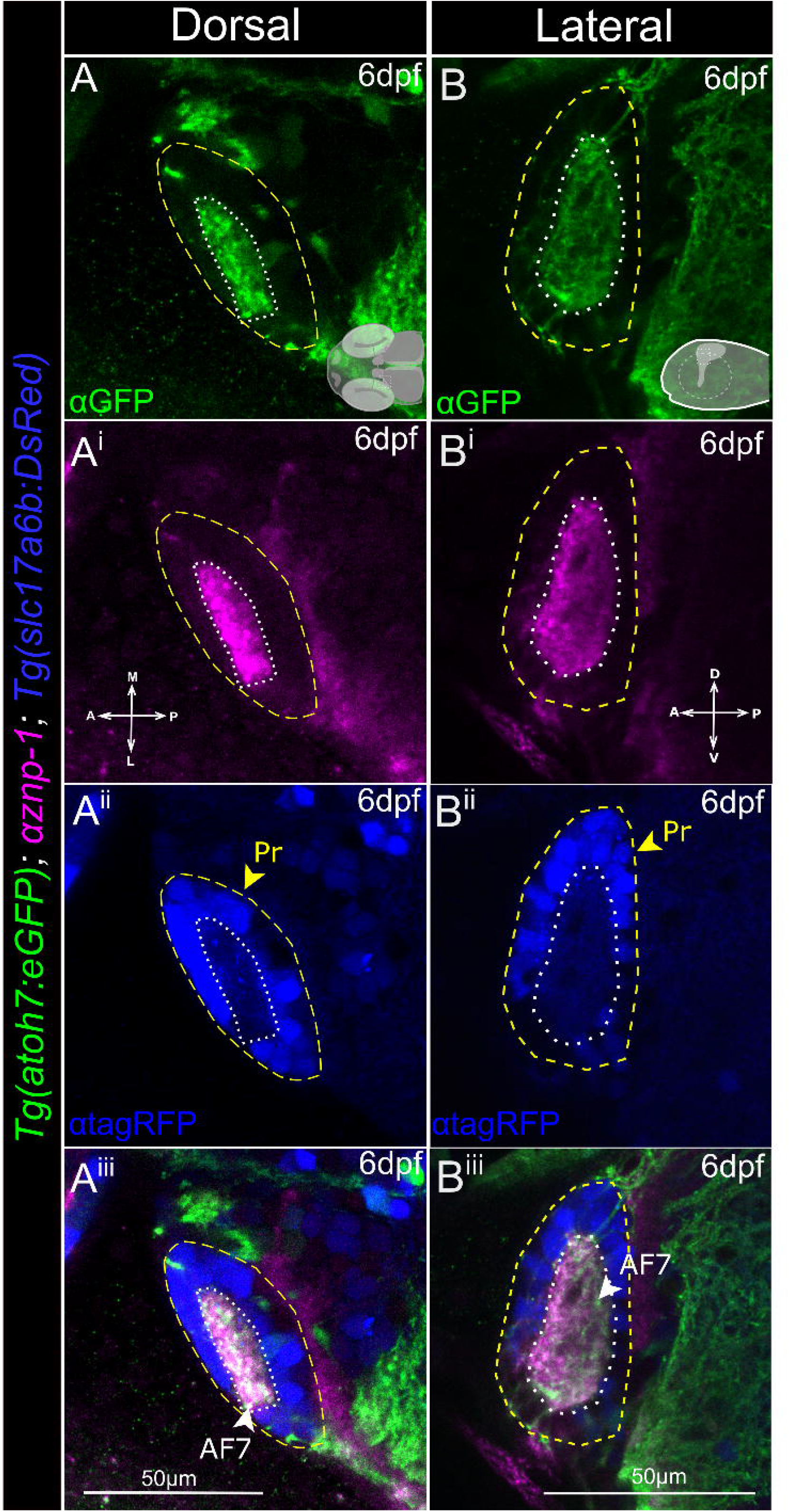
*slc17a6b-positive* neurons surround AF7. **(A-A^iii^)** Dorsal view of partial projection of confocal z-stacks (~5μm). **(B-B^iii^)** Lateral view partial projection (~5μm). *Tg(atoh7:eGFP*) = RGC axons (green), anti-Znp1 = presynaptic terminals (magenta), *Tg(slc17a6b:DsRed*) = glutamatergic neurons (blue), merge = showing AF7 location (green and magenta area). Yellow dashed line outlines Pr region, white dotted line outlines AF7 neuropil. Schematics of orientation are in (A, B).

**SFig. 3.**
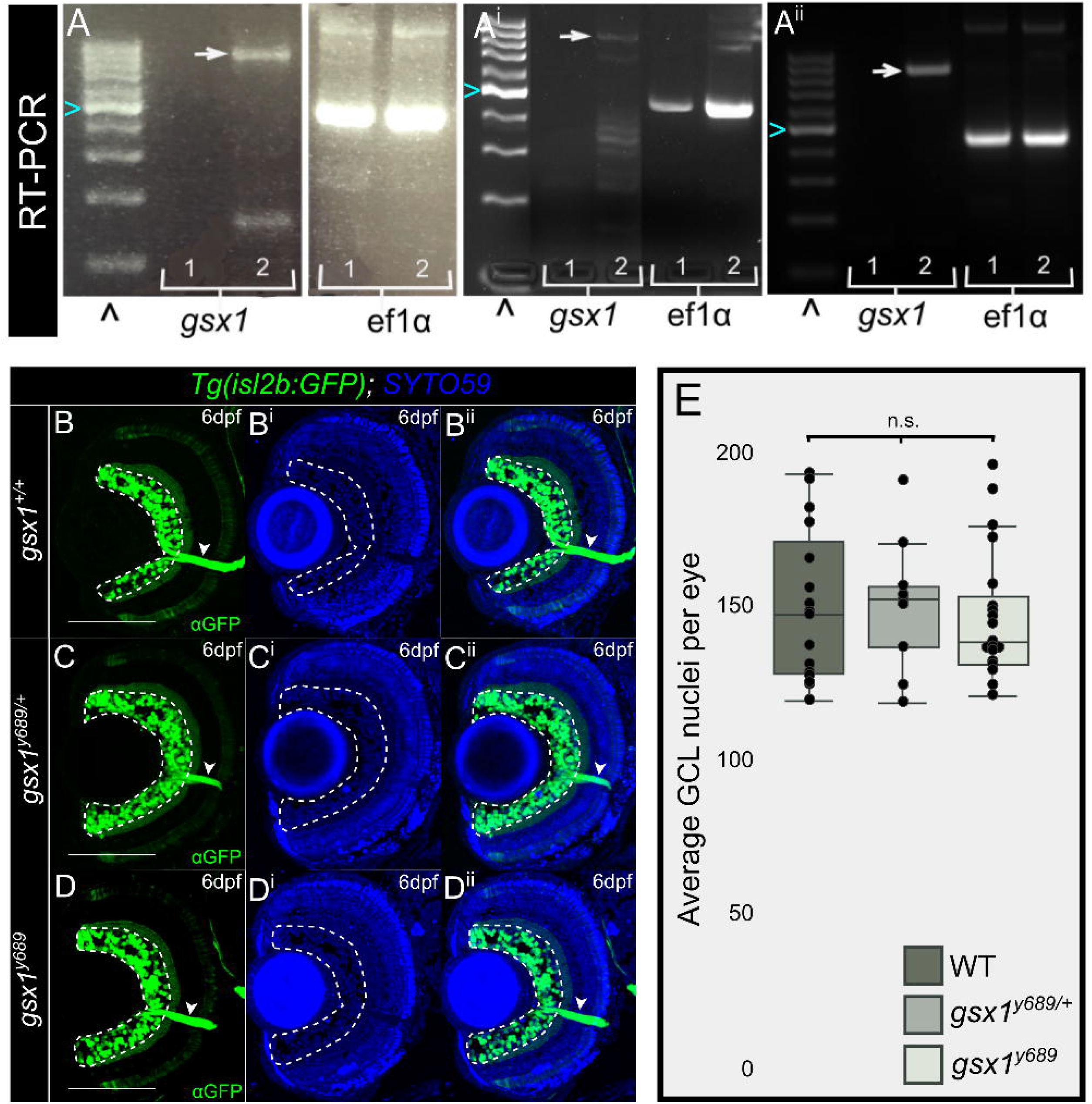
*gsx1* is not expressed in the eye or required for retinal cell type specification or morphology. **(A-A^ii^)** RT-PCR confirming *gsx1* is not expressed in the eye at **(A)** 30 hpf, **(A^i^)** 48 hpf, **(A^ii^)** 6 dpf. 3% agarose gel, white arrow indicates ~800bp *gsx1* fragment. 1 = eye cDNA, 2 = head cDNA. *ef1α* used as a control for DNA quality. ^ = 100bp ladder. Cyan arrowhead = 500bp marker. **(B-D)** Max projections of confocal z-stacks (12μm) of retinal sections at 6 dpf in **(B)** wildtype, **(C)** *gsx1^y689/+^*, **(D)** *gsx1^y689^*. Green = *Tg(isl2b:GFP*), labeling retinal ganglion cells (RGC) in the ganglion cell layer (GCL). Blue = SYTO59, labeling nuclei. White arrow indicates optic nerve leaving the GCL and white dashed outline indicates GCL that is quantified. Scalebar = 100μm. **(E)** Box and whisker plot of GCL quantification for positive RGCs per individual eyes, average taken across 3 consecutive sections with the optic nerve present. One-way ANOVA resulted in no significant differences found across genotypes, *gsx1^+/+^* (n=16), *gsx1^y689/+^* (n=9), *gsx1^y689^* (n=19), F(2,41)=0.11, *p*=0.90.

**SFig. 4.**
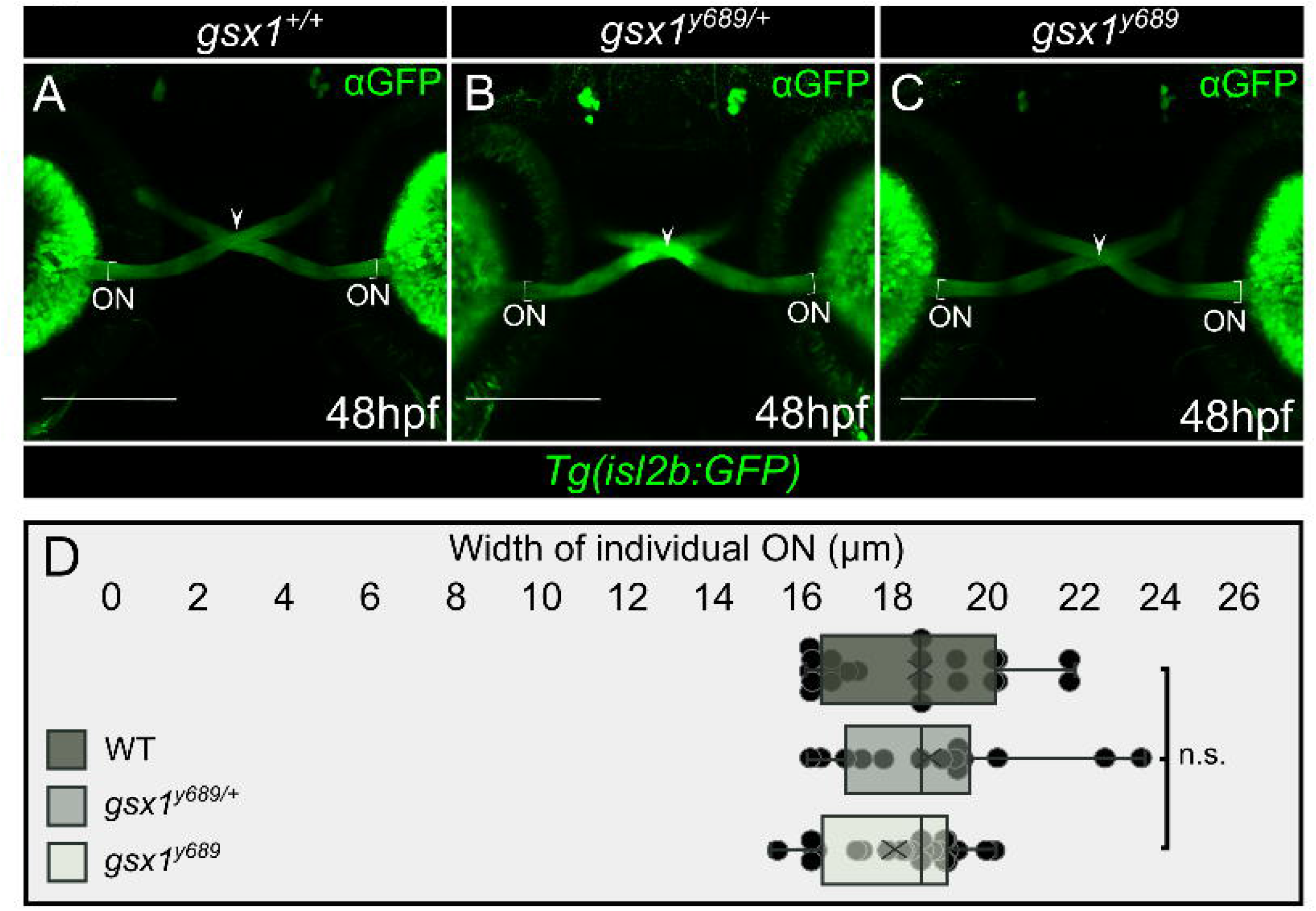
Optic chiasm crossing and optic nerve width are normal in *gsx1* mutants. **(A-C)** Max projection of confocal z-stacks (~55μm) in *Tg(isl2b:GFP*) (green, RGCs) showing normal optic chiasm (white arrowhead) and optic nerve (ON) width at 48 hpf in, **(A)** *gsx1^+/+^* (n=11, 22 ON), **(B)** *gsx1^y689/+^* (n=7, 14 ON), and **(C)** *gsx1^y689^* (n = 12, 24 ON). Bracket outlines optic nerve leaving the eye where measurements were taken. **(D)** Box and whisker plot of measurements for both the right and left ON for each genotype with no statistical differences found, single factor ANOVA [F(2,59)=0.76, *p*=0.47].

**SFig. 5.**
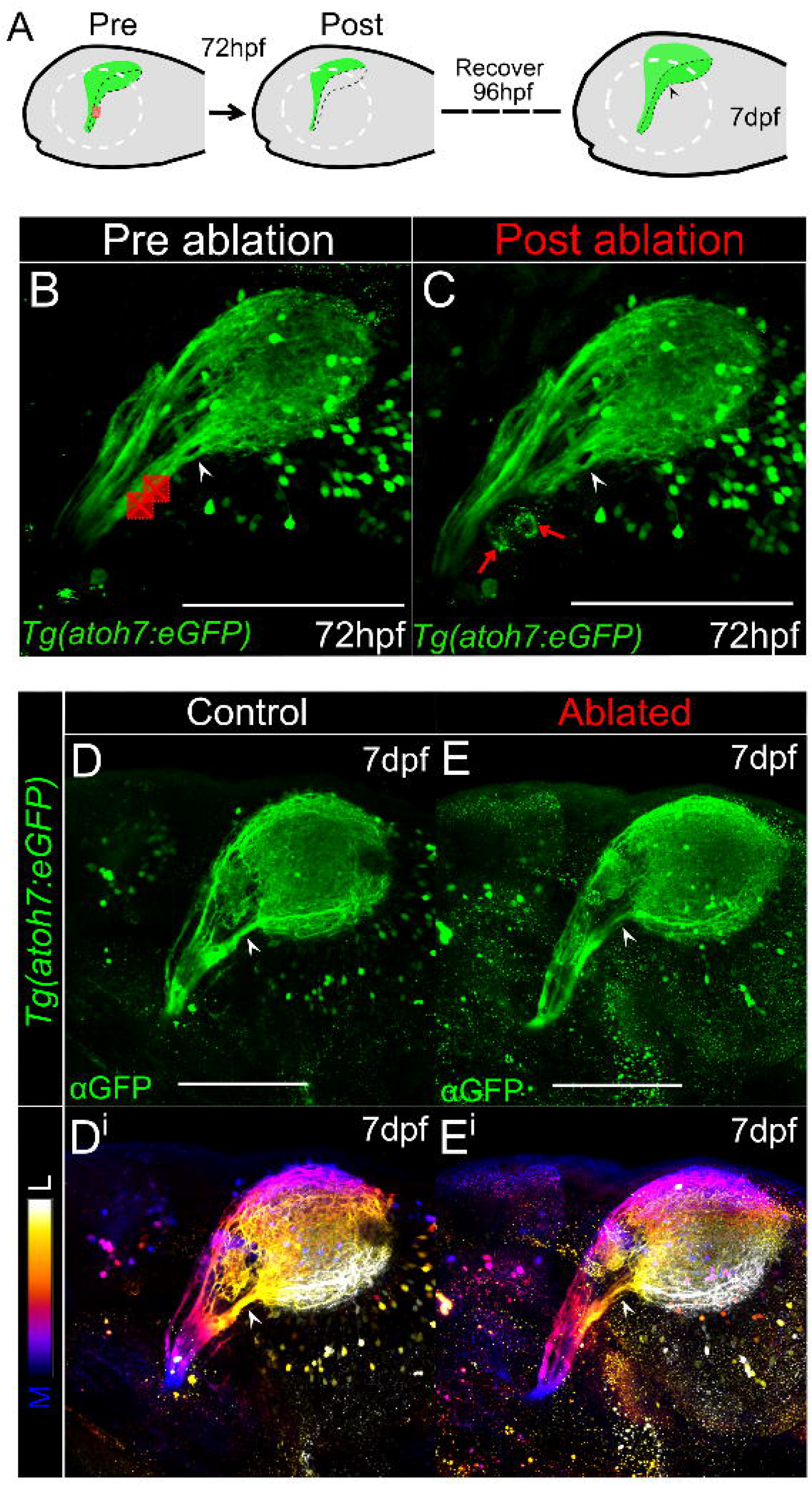
The optic nerve regenerates over the course of four days post 2-photon laser induced injury. **(A)** Experimental timeline and region of targeted ablation following enucleation at 72 hpf. **(B-C)** Lateral orientation of pre and post ablation of max projected 2P z-stacks (~80μm). Red boxes show where ablation took place. Post image red arrows indicate displacement of fluorescent protein following ablation of RGC axons in *Tg(atoh7:eGFP*). Scalebar = 100μm. **(D-E)** 7 dpf max projections of confocal z-stacks in *Tg(atoh7:eGFP*) in **(D)** control non-ablated (n=7) and **(E)** 72 hpf ablated ventral optic nerve (n=8), (~90μm). **(Di, Ei)** RGC axons are depth color coded to provide reference for certain AFs, such as the white arrowhead indicating regeneration of AF6 (yellow). Scalebar = 100μm.

## Notes

### Competing Interest Statement

The authors have declared no competing interest.

